# Dysregulation of mitochondrial function by PLK1-mediated PDHA1 phosphorylation promotes Cr(VI)-associated lung cancer progression

**DOI:** 10.1101/2024.02.16.580630

**Authors:** Qiongsi Zhang, Zhiguo Li, Xiongjian Rao, Derek B. Allison, Qi Qiao, Zhuangzhuang Zhang, Yifan Kong, Yanquan Zhang, Ruixin Wang, Jinghui Liu, Xinyi Wang, Chaohao Li, Fengyi Mao, Wendy Katz, Qing Shao, Tianyan Gao, Xiaoqi Liu

**Affiliations:** Department of Toxicology and Cancer Biology, University of Kentucky, Lexington, Kentucky 40536, USA; Markey Cancer Center, University of Kentucky, Lexington, Kentucky 40536, USA; Department of Pathology and Laboratory Medicine, University of Kentucky, Lexington, Kentucky 40536, USA; Department of Molecular and Cellular Biochemistry, University of Kentucky, Lexington, Kentucky 40536, USA; Department of Chemical and Materials Engineering, University of Kentucky, Lexington, Kentucky 40536, USA; Department of Pharmacology and Nutritional Sciences, University of Kentucky, Lexington, Kentucky 40536, USA

**Author notes:** To whom correspondence should be addressed: Xiaoqi Liu, Department of Toxicology and Cancer Biology, University of Kentucky, Lexington, Kentucky 40536.

## Abstract

Hexavalent chromium (Cr(VI)) is a class I environmental carcinogen known to induce lung epithelial cell transformation and promote lung cancer progression through alterations in the cell cycle and cellular energy metabolism. In this study, we investigated the role of polo-like kinase 1 (PLK1) in Cr(VI)-transformed (CrT) bronchial epithelial cells (BEAS-2B) and found that PLK1 expression was significantly upregulated in CrT cells, leading to impaired mitochondrial function and enhanced cell proliferation both in vitro and in vivo. High levels of PLK1 in CrT cells resulted in decreased mitochondrial activity due to defective modulation of pyruvate dehydrogenase E1 subunit alpha 1 (PDHA1), which is crucial for pyruvate/Acetyl-CoA conversion and carbon influx into the tricarboxylic acid (TCA) cycle. Mechanistically, we demonstrated that PLK1 directly phosphorylates PDHA1 at T57, leading to E1 collapse and PDHA1 degradation via activation of mitophagy. These defects resulted in the inhibition of oxidative phosphorylation and reduction of mitochondrial superoxide generation, ultimately leading to suppression of mitochondrial-mediated apoptotic response. Our findings highlight the role of PLK1 in metabolic reprogramming during Cr(VI)-associated cancer progression, providing new insights and a potential therapeutic target to inhibit Cr(VI)-induced cancer development. Moreover, PLK1 inhibitors may also have the potential to increase chemo-sensitivity of cancer cells by restoring normal mitochondrial function, thereby mitigating drug resistance caused by mitochondrial dysfunction and hyperpolarization.

## Introduction

Hexavalent chromium (Cr(VI)) is not only a well-known carcinogen closely related to the high cancer risk and mortality among workers in various industries using or manufacturing chromate(*1, 2*), but also one of the most common environmental pollutants for the general population. Chromium can be easily assimilated either by inhalation of cigarette smoke(*3*), ambient air(*4*) or ingestion of contaminated drinking-water(*5*). According to the latest case studies from the Center for Disease Control and Prevention (CDC), nearly 558,000 workers in the United States are potentially exposed to chromium and chromium-containing compounds in the workplace. This results in a dramatic increase in the incident rate of lung and bronchus cancer with cumulative exposure(*1, 6*). A recent meta-analysis estimated an overall standardized mortality ratio (SMR) of 1.41 (95%CI: 1.35–1.47) for lung cancer among 47 studies of workers with possible chromium (VI) exposure(*7*). Therefore, the results continuously show that exposure to Cr(VI) significantly associates with the risk of lung and bronchus cancer. Although Cr(VI) has been defined as a human carcinogen, and epidemiology studies also imply an inseparable relationship between Cr(VI) exposure and lung cancer incidence, the mechanisms of Cr(VI)-associated carcinogenesis and tumorigenesis are still undefined. As the molecular mechanism of Cr(VI)-associated carcinogenesis is being explored continuously, it has been confirmed that Cr(VI) promotes chromosomal instability and damage by generation of Cr-DNA adducts(*8*) and DNA oxidation(*9*). Short-period exposure to Cr(VI) leads to DNA damage-mediated cell cycle arrest and apoptosis(*10*), whereas chronic exposure induces cell morphology changes, cell transformation and neoplastic transformation(*11, 12*). The establishment of cell-cycle checkpoints in response to chromium exposure-mediated genotoxic stress is essential for genome integrity. However, bypass of checkpoints enhances Cr (VI)-associated mutagenesis, followed by the acquisition of certain genetic or epigenetic modifications that give cells a stronger growth advantage and result in uncontrolled proliferation. Studies reported by Lal et al revealed that Akt1 activation can bypass the G1/S checkpoint induced by Cr(VI) exposure in normal human diploid cells(*13*). In following studies, Chun et al reported that another protein serine/threonine kinase, PLK1, is necessary for bypass of the G_2_/M checkpoint after Cr(VI) exposure in HLF cells and in S.cerevisiae(*14*). In addition to the growth advantages caused by bypass of checkpoints in the face of Cr(VI) exposure, chronic Cr(VI) exposure results in metabolic reprogramming, both in cultured cells and in isolated mitochondria(*15*). Reprogrammed metabolism is one of the hallmarks of cancer cells. Recent studies with human normal bronchial epithelial BEAS-2B cells, prostate carcinoma DU145 cells and isolated rat liver mitochondria further confirmed the previous results, as they showed that both short-term treatment and chronic exposure to Cr(VI) induced a shift in energy metabolism(*16-18*). Cr(VI)-associated cancer progression not only requires cells to overcome the growth limitation imposed by cell cycle regulation, but also requires a supply of energy sufficient to provide growth momentum, and requires cells to adapt to excessive apoptotic stress caused by DNA damage and reactive oxygen species (ROS). These are established in the long-term adaptation of cells to Cr(VI) treatment. During the dynamic adaptation process, transformed cells gradually gain advantages in division and survival. The pivotal mechanisms that play a tandem role in chromium-associated cell transformation and cancer progression remain unknown and require extensive investigation.

PLK1, a well-established mitotic regulator, not only plays a pivotal role in cell-cycle regulation but is also involved in a diverse range of biological events. Dysregulation of PLK1 has been shown to disrupt many of these events (*19*). In addition, PLK1 overexpression has been reported in a variety of cancers including lung cancer(*20-22*), and higher PLK1 expression has been correlated with poor prognosis of cancer(*23, 24*). Pre-clinical evidence suggests that the molecular targeting of PLK1 could be an effective therapeutic strategy in a wide range of cancers(*25*). Among the PLK1-related mechanisms that promote cancer progression, aspects of the regulation of cancer cell metabolism have been frequently mentioned. Of note, numerous reports demonstrated a potential relationship between PLK1 and reprogrammed cancer cell metabolism. Furthermore, tumor cell growth was inhibited by regulating certain core metabolic pathways with PLK1 inhibitors. Among them, Cholewa and colleagues reported that PLK1 inhibition with the small-molecule ATP-competitive PLK1 inhibitor BI6727 (Volasertib) resulted in a decrease in multiple metabolic proteins (*26*). In our previous studies, we also demonstrated that PLK1 phosphorylation of phosphatase and tensin homolog (PTEN) causes a tumor-promoting metabolic state, promoting the Warburg effect(*27*). Although it was indicated that PLK1 also has a potential regulatory role in oxidative phosphorylation (OXPHOS) (*28, 29*), details of the underlying mechanisms are still lacking. Compared with normal cells, energy metabolism in cancer cells is dramatically altered, and the role of PLK1 in this process remains to be elucidated. A more thorough study of PLK1 will enable us to comprehensively understand the accumulation of growth advantages, the transformation of energy metabolism, immune evasion and anti-apoptosis of cancer cells. This will provide a new target for cancer therapy, thereby limiting the occurrence and progression of cancer.

In our investigation into the diverse functions of PLK1 in types of cancer and their progression, we found that the level of PLK1 was significantly increased, and mitochondrial activity was decreased along with metabolic reprogramming during Cr(VI)-mediated cell transformation. Subsequent manipulation of PLK1 protein level and kinase activity rescued mitochondrial activity and resulted in a substantial reduction in cancer cell proliferation in both in vitro and in vivo settings. Mechanistically, our finding unveiled that PDHA1, a key enzyme for oxidative phosphorylation, is involved in the PLK1-mediated metabolic reprogramming. Specifically, we demonstrated that PLK1 phosphorylation of PDHA1 at T57 triggers a collapse in E1 heterotetramer binding, which results in downregulation of oxidative phosphorylation, inhibition of mitochondrial activity and eventually activates mitophagy. We also performed stable isotope-resolved metabolomics to reveal the detailed carbon influx from glucose to TCA cycle, which emphasizes the fact that PLK1 is tightly involved in glucose metabolism and negatively regulates TCA cycle through phosphorylation of PDHA1 at T57. Furthermore, we extended our investigation to the metabolism of whole mice, using a mouse model with mutated Pdha1 expressed (Pdha1 T57D conditional knock-in). We found that mice with Pdha1 T57D knock-in had a lower metabolic rate and oxygen consumption compared to the control mice. Collectively, our comprehensive findings provide compelling evidence for the potential of combining PLK1 inhibition with the use of the PDH activator dichloroacetate (DCA) as a novel therapeutic approach for the treatment of Cr(VI)-associated lung cancer. These findings highlight the therapeutic implications of our research in the development of innovative strategies for lung cancer treatment.

## Results

### Expression of PLK1 is elevated during Cr(VI)-associated cell transformation, and knockdown of PLK1 inhibits Cr(VI)-mediated cancer progression

Cr(VI)-associated cell transformation enhances cell proliferation and interferes cell cycle regulation. As one of the key regulators of the cell cycle, PLK1 functions in eukaryotic cell division. It has been reported that PLK1 is closely related to cancer progression and is highly expressed in multiple cancer types. To directly ask whether PLK1 is involved in Cr(VI)-associated carcinogenesis, we analyzed the protein levels of PLK1 in human bronchial epithelial cells (BEAS-2B) and Cr(VI)-transformed human bronchial epithelial cells (CrT). Compared to BEAS-2B cells, CrT cells have a much higher level of PLK1 protein, indicating that Cr(VI)-associated transformation results in PLK1 overexpression (Figure 1A). In addition to examining transformed cells due to chronic exposure to Cr(VI), we also performed a short-term sodium dichromate treatment of BEAS-2B cells. With increasing concentrations of sodium dichromate, the expression levels of PLK1 were upregulated accordingly (Figure 1B). Therefore, we conclude that both long-term treatment with Cr(VI) and short-term Cr(VI) exposure result in elevation of PLK1 in BEAS-2B cells.

**Figure 1.**
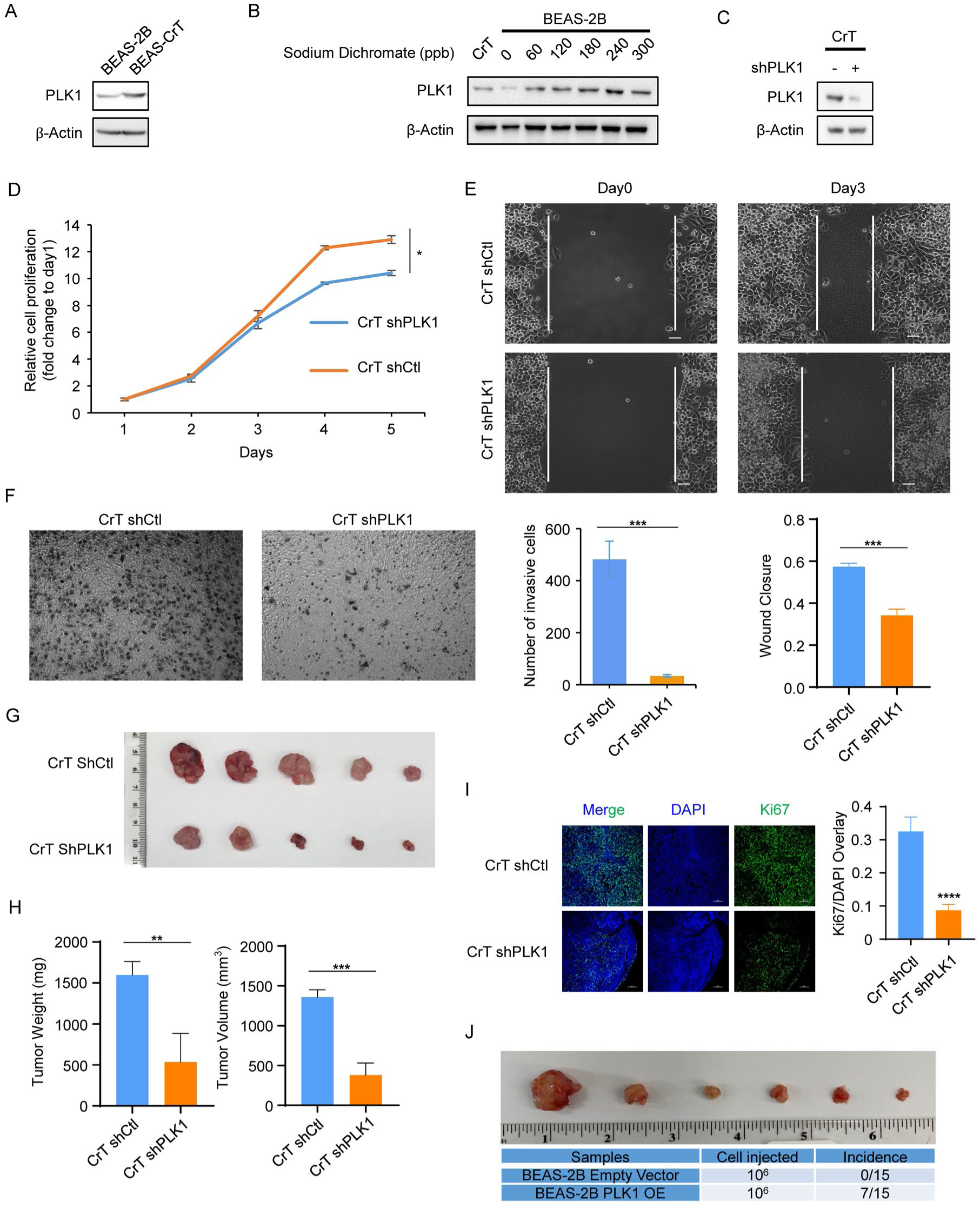
Expression of PLK1 is upregulated during Cr(VI)-mediated cell transformation, and knockdown of PLK1 inhibits Cr(VI)-associated cancer progression. (A) Immunoblotting (IB) analysis of PLK1 protein level in BEAS-2B and BEAS-2B Cr(VI) transformed (CrT) cells. (B) BEAS-2B cells were treated with sodium dichromate at the indicated doses (60, 120, 180, 240 and 300 ppb) for 12h, and harvested for IB. CrT cells were used to indicate the level of PLK1 after chronic Cr(VI) exposure. (C) IB of PLK1 protein level in CrT cells stably expressing a control shRNA or PLK1 knockdown shRNA. (D) MTT analysis of CrT cells stably expressing a control shRNA or PLK1 knockdown shRNA. The results are presented as mean ± standard deviations (n=6), *P<0.05. (E) Representative images of wound healing analysis in day 0 and day 3 to compare the cell migration rates in CrT cells stably expressing a control shRNA or PLK1 knockdown shRNA. Quantification of wound healing analysis in CrT shCtl and CrT shPLK1 cells and the results are presented as means ± standard deviations (n=3). ***P<0.001. (F) Representative images of cell invasion analysis to compare the cell invasion rates in CrT shCtl and CrT shPLK1 cells. Results are presented as mean ± standard deviations (n=3). ***P<0.001. (G and H) CrT shCtl and CrT shPLK1 cells were inoculated into flanks of nude mice (n=10), tumor images, weights and volumes are presented. **P<0.01; ***P<0.001. (I) IF staining for Ki67, a marker for cell proliferation, shows fewer proliferating cells in tumors with PLK1 knockdown. Quantification of Ki67 staining is presented. ****P<0.0001. (J) Parental BEAS-2B cells with empty vector and BEAS-2B cells stably overexpressing PLK1 were inoculated into flanks of nude mice (n=15). Images of tumors and tumor incidence.

To examine the significance of PLK1 in Cr(VI)-associated carcinogenesis, we performed PLK1 knockdown in CrT cells by using shRNA, followed by anti-PLK1 IB to verify the knockdown efficiency (Figure 1C). Furthermore, CrT cells with PLK1 knockdown showed a reduced level of cell proliferation than that of parental cells (Figure 1D). Moreover, compared to the parental cells, CrT cells with stable knockdown of PLK1 also showed a significant decrease in cell migration (Figure 1E) and cell invasion ability (Figure 1F). These results together indicate that PLK1 plays a positive role in Cr(VI)-associated carcinogenesis.

To determine the proliferation rate of PLK1-knockdown cells in vivo, equal numbers of CrT shCtl and CrT shPLK1 cells were subcutaneously injected into nude mice. Both the tumor weights and tumor volumes in the PLK1 knockdown group were reduced (Figures 1G & 1H). Moreover, PLK1 knockdown resulted in significant inhibition of tumor cell proliferation as indicated by IF staining against Ki67 (Figure 1I). Histological examination of xenograft tumors derived from CrT cells also showed that the number of mitotic cells was significantly lower in shPLK1 tumors compared to shCtl tumors (Figure S1B). Thus, downregulation of PLK1 inhibits cancer cell proliferation in vivo. Furthermore, two groups of BEAS-2B cells were transduced with lentivirus encoding PLK1 and empty vector. Equal numbers of BEAS-2B control (BEAS-2B E.V) cells and BEAS-2B PLK1-overexpression (BEAS-2B PLK1-OE) cells were injected subcutaneously into nude mice. Tumor incidence and tumor volumes for mice carrying BEAS-2B PLK1-OE cells were dramatically increased compared to mice with BEAS-2B Ctl cells (Figures 1J and Figure S1A). In conclusion, PLK1 plays a pivotal role in promoting Cr(VI)-associated carcinogenic properties, enhancing Cr(VI)-associated cancer progression both in vitro and in vivo.

### PLK1 elevation results in metabolic reprogramming

Several prior studies reported that Cr(VI)-associated cell transformation results in metabolic reprogramming. Moreover, Cr(VI) exposure can cause respiration inhibition and defective mitochondrial function in vivo, ex vivo, in isolated mitochondria and in intact cells. In addition, increasing evidence suggests that PLk1 activation is linked to tumor-promoting metabolic status. Because PLK1 promotes a tumor-promoting metabolic status, i. e., increased glycolysis, we speculated that PLK1 may be critical for regulating mitochondrial activity and oxidative phosphorylation in Cr(VI)-associated cancer progression. To verify our hypothesis, we treated CrT cells with the PLK1 inhibitor BI6727, followed by a mitochondrial stress test using Seahorse Bioscience XFe96 analyzer. Inhibition of PLK1 kinase activity indeed enhanced basal respiration and spare respiratory capacity in CrT cells, supporting the notion that PLK1 plays a significant role to regulate OXPHOS. We extended similar test to H2009 cells, a lung adenocarcinoma cell line, which confirmed our finding that inhibition of PLK1 kinase activity enhances OXPHOS (Figure 2A). To further determine whether PLK1 is involved in the regulation of OXPHOS in Cr(VI)-associated cell transformation, we compared the basal respiration and spare respiratory capacity in CrT shCtl and CrT shPLK1 cells. Although PLK1 knocking down showed limited effects on basal respiration, CrT shPLK1 cells had a higher level of spare respiratory capacity compared to CrT shCtl cells, indicating that cells with a lower protein level of PLK1 may have a higher mitochondrial activity level and a higher capability to generate extra ATP through oxidative phosphorylation under stress conditions. (Figure 2B). In support of this, PLK1 downregulation also enhanced mitochondrial activity and rescued mitochondrial membrane potential in CrT cells (Figure S2A). We also performed a mitochondrial stress test in BEAS-2B cells and BEAS-2B PLK1-OE cells. In agreement with the results based on PLK1 knockdown, we show that BEAS-2B cells with PLK1 overexpression tended to have a lower level of both basal respiration and spare respiratory capacity (Figure 2C). Thus, we conclude that PLK1 acts as a negative regulator of oxidative phosphorylation and mitochondrial activity.

**Figure 2.**
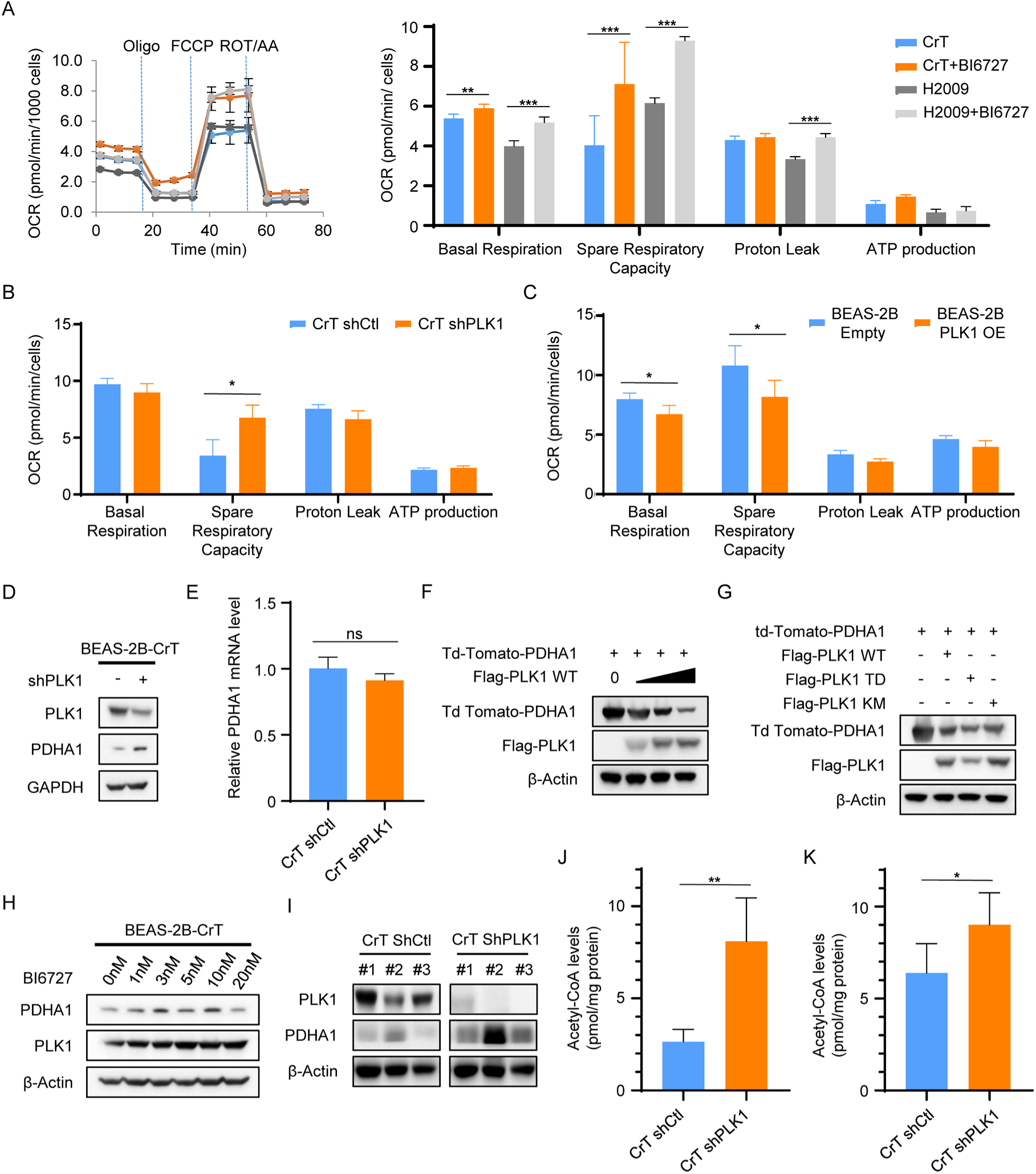
PLK1 elevation results in metabolic reprogramming. (A, B and C) Representative traces of the oxygen consumption rates (OCR), when oligomycin (Oligo), carbonyl cyanide-4- (trifluoromethoxy) phenylhydrazone (FCCP), rotenone plus antimycin A (ROT/AA) were injected into the assay XF96 plates. Each data point is mean ± standard deviation (n=6). Quantification of metabolic parameters are presented as mean ± standard deviations. *P<0.05; **P<0.01; ***P<0.001. The mitochondrial stress analyses were performed in (A) CrT and H2009 cells treated with or without 25nM BI6727 for 16h; (B) CrT shCtl and CrT shPLK1 cells; (C) BEAS-2B cells with empty vector and BEAS-2B cells with PLK1 overexpression. (D) IB of PLK1 and PDHA1 in CrT cells and CrT cells depleted of PLK1 with shRNA. (E) qRT-PCR analysis of transcriptional levels of PDHA1 in CrT shCtl and CrT shPLK1 cells. (F) HEK293TA cells were co-transfected with tdTomato-PDHA1 and increasing levels of Flag-PLK1, followed by IB analysis of tdTomato-PDHA1 and Flag-PLK1. (G) HEK293TA cells were co-transfected with tdTomato-PDHA1 and different forms of Flag-PLK1 (WT, -T210D and -K82M mutant), followed by IB. (H) CrT cells were treated with increasing doses of BI6727 for 16h. (I) IB analysis of PLK1 and PDHA1 in lysates collected from CrT shCtl and CrT shPLK1-derived xenografts. (J) Acetyl-CoA levels in CrT shCtl and CrT shPLK1 cells are presented as means ± standard deviations (n=3). **P<0.01. (K) The levels of acetyl-CoA in samples from CrT shCtl and CrT shPLK1 cell-derived xenografts are presented as mean ± standard deviations (n=3). *P<0.05.

Having established that PLK1 is a negative regulator of OXPHOS, we examined the underlying mechanism for this novel observation. We focused our attention to PDHA1, a critical molecule that links glycolysis and OXPHOS. As indicated in Figure 2D, knock-down of PLK1 in CrT cells led an upregulation of the protein level of PDHA1. Furthermore, a doxycycline (DOX)-inducible PLK1-knockdown H2030 cell line was generated and utilized to further test the effect of PLK1 on PDHA1. The results showed that the protein level of PDHA1 was significantly increased with the knockdown of PLK1 induced by DOX treatment (Figure S2B). Pyruvate serves as a bridge between glycolysis in the cytosol and oxidative phosphorylation in mitochondria. The pyruvate dehydrogenase complex catalyzes conversion from pyruvate to acetyl-CoA, which drives the tricarboxylic acid (TCA) cycle. Therefore, we tested the hypothesis that PLK1-associated metabolic reprogramming might rely on the PLK1-PDHA1 regulatory axis. Accordingly, CrT shCtl and CrT shPLK1 cells were collected, followed by measurement of the levels of PDHA1 protein and mRNA. As an important functional protein in mitochondria, the function and protein level of PDHA1 are regulated by multiple mechanisms including regulation of transcription, post-translational modification and mitochondrial biogenesis. To first rule out a potential effect of PLK1 on mitochondrial biogenesis, we demonstrated that the transcriptional level of mtDNA-encoded cytochrome b (mtCYB) was not significantly altered in CrT shPLK1 cells compared with either CrT shCtl cells or parental BEAS-2B cells (Figure S2C). Notably, PLK1 knockdown only resulted in upregulation of the PDHA1 protein level (Figure 2D), but not PDHA1 transcriptional level (Figure 2E), indicating that PLK1-associated PDHA1 downregulation was likely due to post-translational modification. In other words, PLK1 may induce PDHA1 protein degradation. To directly address whether PLK1 triggers PDHA1 protein degradation, we co-expressed tdTomato-PDHA1 and Flag-PLK1 in HEK293TA cells. As shown in Figure 2F, PLK1 downregulated the protein level of PDHA1 in a PLK1-dosage dependent manner. To further confirm whether the kinase activity of PLK1 was required for the degradation of PDHA1, we overexpressed the same amount of PDHA1 as well as 3 different PLK1 constructs (wild type, the constitutively active T210D variant, and the kinase-defective K82M variant) and we found that expression of the T210D variant led to the lowest PDHA1 protein level (Figure 2G). In support, treatment of CrT cells with PLK1 inhibitor BI6727 also led to an increased level of PDHA1, suggesting that endogenous PDHA1 is inhibited by endogenous PLK1 in a kinase activity-dependent manner (Figure 2H). Furthermore, PDHA1 protein level was increased in xenograft tumors upon PLK depletion (Figure 2I). To support the concept that PLK1 is a negative regulator of PDHA1, we showed that the levels of acetyl-CoA, the final catalytic product of PDH, were significantly upregulated in both CrT cells (Figure 2J) and xenografts derived from CrT cells upon PLK1 knock-down (Figure 2K). In summary, PLK1-associated inhibition of OXPHOS is due to downregulation of PDHA1 both in vitro and in vivo.

### PLK1-mediated phosphorylation of PDHA1 enhances its degradation

Having established that PLK1 is a negative regulator of PDHA1, we dissected the molecular details of the PLK1/PDHA1 axis. We first asked whether PDHA1 directly interacts with PLK1. Accordingly, HEK293TA cells were transfected with Flag-PLK1 as well as tdTomato-PDHA1, and harvested for anti-Flag IP, followed by anti-tdTomato IB. Meanwhile, HEK293TA cells expressing Flag-PLK1 and tdTomato-PDHA1 were harvested for anti-tdTomato IP, followed by anti-Flag IB. We consistently found that ectopically expressed PDHA1 and PLK1 bound to each other in both co-IP assays (Figures 3A, 3B). Next, we examined whether endogenous PDHA1 binds to PLK1. For that purpose, CrT cells were harvested for anti-PLK1 IP, followed by anti-PDHA1 IB. Similar to binding between exogenous PLK1 and PDHA1, we also found that endogenous PDHA1 directly binds to PLK1 in CrT cells (Figure 3C). To address whether PDHA1 is a direct substrate of PLK1, a series of in vitro kinase assays were performed with purified PLK1 and recombinant PDHA1 in the presence of [γ-^32^P] ATP. As indicated in Figure 3D, we showed that PLK1 phosphorylates PDHA1 in vitro. To map the phosphorylation site(s), we mutated all the potential serine and threonine residues into alanine, and identified T57 as the PLK1 phosphorylation site (Figure 3E). To further verify this phosphorylation site, a polyclonal antibody that specifically targets the phosphorylated form of PDHA1 at T57 was generated (Figure 3F).

**Figure 3.**
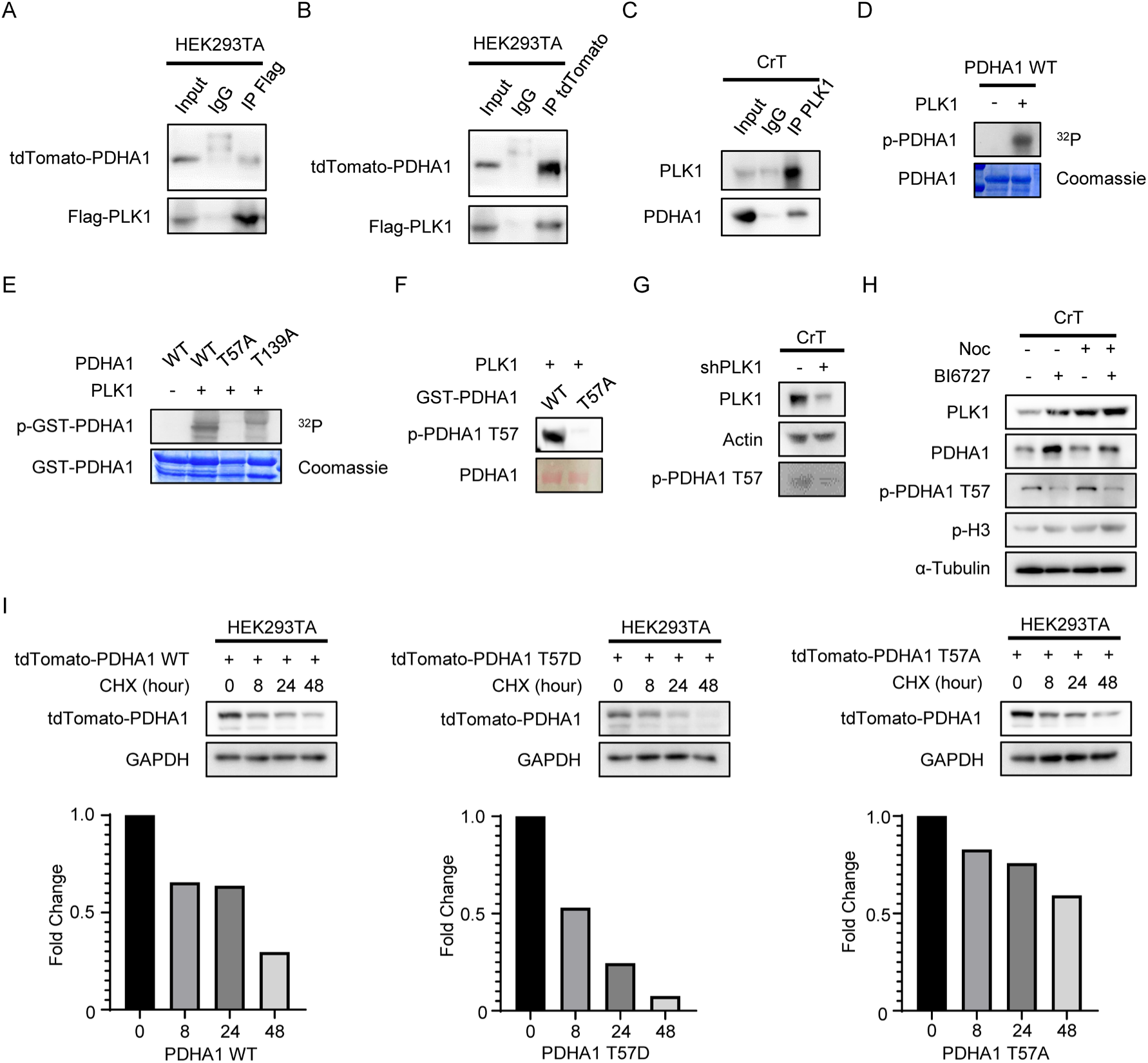
PLK1-mediated PDHA1 phosphorylation enhances its protein degradation. (A) HEK293TA cells were co-transfected with Flag-PLK1 and tdTomato-PDHA1, and harvested for anti-Flag immunoprecipitation (IP), followed by IB to detect tdTomato-PDHA1 and Flag-PLK1. (B) HEK293TA cells were co-transfected with Flag-PLK1 and tdTomato-PDHA1, and harvested for anti-tdTomato IP, followed by IB to detect tdTomato-PDHA1 and Flag-PLK1. (C) Lysates of CrT cells were subjected to anti-PLK1 IP, followed by IB to detect PLK1 and PDHA1. (D) Purified PLK1 was incubated with purified PDHA1 in the presence of [γ-^32^P] ATP. The reaction mixtures were resolved by SDS-PAGE, stained with Coomassie brilliant blue, and detected by autoradiography. (E) PLK1 was incubated with recombinant GST-PDHA1 (WT, T57A or T139A) as in D. (F) Recombinant GST-PDHA1 (WT or T57A) were incubated with purified PLK1 in the presence of ATP, followed by IB against phospho-T57-PDHA1. Bottom panel shows the protein level by ponceau S staining. (G) IB analysis of PLK1 and pT57-PDHA1 in CrT shCtl and CrT shPLK1 cells. (H) CrT cells were treated with 100ng/mL nocodazole and 10nM BI6727 for 16h, and harvested for IB against PLK1, PDHA1 and pT57-PDHA1. (I) HEK293TA cells were transfected with different forms of tdTomato-PDHA1 constructs (WT, T57D and T57A), treated with 20μg/ml cycloheximide (CHX) for the indicated times, and harvested for IB. Normalized fold change is presented.

With the pT57-PDHA1 antibody, we addressed whether endogenous PDHA1 can be phosphorylated by endogenous PLK1 at T57 in vivo. As indicated, the pT57-PDHA1 antibody was able to detect the phosphorylation signal of PDHA1 in CrT cells, but the epitope was dramatically reduced in CrT cells upon PLK1 knockdown (Figure 3G). Similarly, doxycycline-induced PLK1 knock-down in H2030 cells decreased the phosphorylation level of PDHA1 at T57 (Figure S2B). Requirement of PLK1-associated kinase activity for pT57-PDHA1 epitope was further tested with the PLK1 inhibitor BI6727. Accordingly, CrT cells were treated with nocodazole or BI6727 and harvested for IB. As shown in Figure 3H, nocodazole treatment increased the level of pT57-PDHA1 and co-treatment with BI6727 clearly inhibited the level of pT57-PDHA1, supporting the notion that the PDHA1-T57 phosphorylation is PLK1 kinase activity dependent. Although nocodazole-associated mitotic arrest and PLK1 elevation resulted in elevation of pT57-PDHA1 phosphorylation, the level of PDHA1 protein was clearly decreased, indicating that PLK1-dependent phosphorylation of PDHA1 at T57 drives its degradation. To test this hypothesis directly, we compared the protein turnover of different forms of PDHA1 (WT, phospho-mimetic T57D variant and PLK1-unphosphorylatable T57A variant) with the cycloheximide (CHX) chasing assay. Compared to WT-PDHA1, the protein level of PDHA1-T57D variant was quickly diminished upon inhibition of protein synthesis. In contrast, turnover of PDHA1 protein was markedly inhibited in the cells expressing PDHA1-T57A variant (Figure 3I). In summary, we concluded that PLK1-mediated PDHA1-T57 phosphorylation drives its degradation.

### PLK1-associated PDHA1 phosphorylation triggers activation of mitophagy and promotes PDHA1 protein degradation

Having established that PLK1 phosphorylates PDHA1 at T57 and enforces PDHA1 protein degradation, we then explored the potential mechanisms that are responsible for the phospho-T57-mediated PDHA1 degradation. Eukaryotic cells have two major protein degradation pathways: the ubiquitin proteasome system (UPS) and the autophagy-lysosomal pathway (ALP). To characterize the pertinent regulatory pathway, 293TA cells transfected with tdTomato-PDHA1 and Flag-PLK1 were treated with or without the proteasome inhibitor MG132 (25µM) for 4 hours. As shown in Figure 4A, the expression level of PDHA1 was downregulated upon PLK1 co-expression compared to control. However, turnover of PDHA1 protein failed to be rescued with inhibition of proteasome activity. Thus, PLK1-associated PDHA1 protein degradation is not due to proteasomal degradation. To further characterize the degradative mechanisms, 293TA cells transfected with tdTomato-PDHA1 and Flag-PLK1 were treated with or without the autophagy inhibitor 3-MA (3mM) for 4 hours. Surprisingly, PDHA1 protein, which was attenuated by PLK1 co-expression, returned to the control level upon inhibition of autophagy, suggesting that PLK1-associated PDHA1 protein degradation depends on autophagy (Figure 4B). To further confirm this conclusion, 293TA cells transfected with two tdTomato-PDHA1 constructs (WT and T57A) were treated with 3 different autophagy/mitophagy inhibitors, 3-MA, chloroquine and liensinine following 100ng/mL nocodazole treatment. We found that wild type PDHA1 protein degradation was enhanced by nocodazole treatment and blocked by autophagy/mitophagy inhibition. For cells expressing the T57A variant of PDHA1, neither nocodazole nor autophagy/autophagy inhibitor treatment affected the PDHA1 protein level. Therefore, PDHA1 protein degradation in PLK1 highly upregulated cells is mediated through autophagy, and elimination of PDHA1-T57 phosphorylation prevents autophagy/mitophagy-mediated PDHA1 degradation (Figure 4C). In sum, we conclude that PLK1-associated PDHA1 degradation depends on ALP rather than UPS.

**Figure 4.**
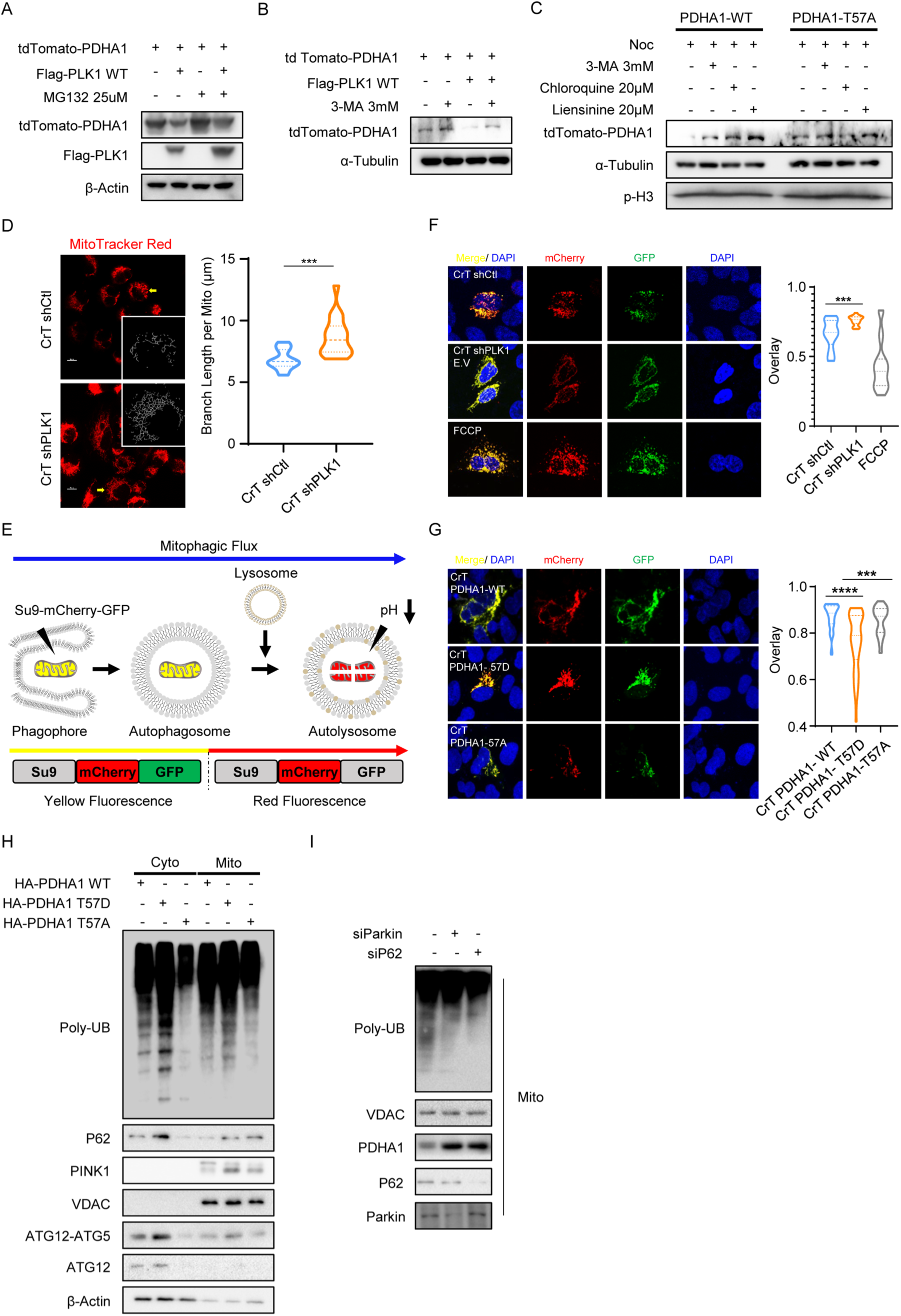
PLK1-associated PDHA1 phosphorylation triggers mitophagy activation and promotes PDHA1 protein degradation. (A) HEK293TA cells were co-transfected with tdTomato-PDHA1 and Flag-PLK1, treated with 25μM MG132, and harvested for IB. (B) HEK293TA cells were co-transfected with tdTomato-PDHA1 and Flag-PLK1, treated with 3mM 3-MA, and harvested for IB. (C) HEK293TA cells were transfected with tdTomato-PDHA1 (WT and T57A), treated with 100ng/ml nocodazole overnight, incubated with 3-MA, chloroquine and liensinine as indicated, and harvested for IB. (D) CrT shCtl and CrT shPLK1 cells were stained with MitoTracker (red), followed by confocal microscopy analysis, with skeletonized mitochondria being presented as inserts. After single cells were analyzed using ImageJ plug Marcros, quantitative analysis and comparison of mitochondrial mean branch length between CrT shCtl cells and CrT shPLK1 cells are presented on the right panel. ***P<0.001. (E) The scheme illustrates the working mechanism of the mitophagy flux indicator. The mitophagy activation index was calculated based on the degree of GFP and mCherry co-localization as measured by Pearson coefficient. Decreased co-localization (i.e. a decrease in mitophagy index) indicates increased mitophagy in cells. (F) CrT cells (shCtl or shPLK1) were transfected with the plasmid expressing the mitophagy indicator for 2 days, and analyzed by the confocal microscopy, followed by quantification of the mitophagy activation level. ***P<0.001. (G) CrT cells stably expressing different forms of PDHA1 constructs (WT, T57D and T57A) were transfected with the plasmid expressing mitophagy indicator for 2 days, and analyzed under the confocal microscope, followed by quantification to measure the mitophagy activation. ***P<0.001; ****P<0.0001. (H) CrT cells stably expressing wild type PDHA1 and PDHA1 variants (T57D and T57A) were harvested for subcellular fractionation, followed by IB to address the level of poly-ubiquitination (Poly-UB), P62, PINK1, ATG12 and ATG12-ATG5 conjugation. (I) Parkin and P62 knockdown by siRNA in CrT cells, followed by mitochondrial isolation and IB.

When we further explored whether autophagy or mitophagy underly PLK1-associated PDHA1 degradation, we noticed that BEAS-2B cells have a more complete mitochondrial network compared to CrT cells. Confocal microscopy was used to characterize the mitochondrial morphology of both BEAS-2B and CrT cells. Figure S3A illustrates qualitatively how elongation of the mitochondria was severely undergone mitochondrial fission during cell transformation from a filamentous phenotype observed in the BEAS-2B cells, to appearing fragmented and aggregated perinuclearly in the CrT cells. To further confirm PLK1-associated mitochondrial morphology change, similar experiments were performed in CrT shCtl cells and CrT shPLK1 cells. Interestingly, CrT shPLK1 cells showed a more complete mitochondrial network compared to CrT shCtl cells. As shown in Figure 4D, we observed an increased mitochondrial mean branch length in CrT shPLK1 cells compared to CrT shCtl cells, suggesting that PLK1 triggers mitochondrial fission and that downregulation of PLK1 restores the integrity of the mitochondrial network. In addition, we noticed that PLK1 phosphorylation of PDHA1 at T57 results in PDHA1 re-localization from mitochondria to cytosol in BEAS-2B cells (Figures S3B and S3C). Further, we demonstrated that PDHA1-T57D variant was localized in both cytosol and mitochondria in CrT cells, whereas PDHA1-T57A and wild type PDHA1 located exclusively in mitochondria (Figure S3D). Mitochondria are highly dynamic organelles that undergo continuous fission, fusion, biogenesis and mitophagy, all of which maintain mitochondrial morphology, homeostasis and inheritance in different cases of bioenergetics or oxidative challenges. Given that PLK1 is highly involved in mitochondrial dysfunction and reforming, and that PLK1 triggers PDHA1 protein degradation in an ALP-dependent manner, we assume that PLK1-associated PDHA1 degradation is likely mediated through mitophagy, which results in change of mitochondrial morphology.

To provide more direct evidence that mitophagy is involved in PLK1-associated PDHA1 degradation in CrT cells, we delivered the plasmid expressing a mitophagy indicator (matrix-targeted Su9-mCherry-GFP)(*30*) into the cells. Su9-mCherry-GFP is first expressed in mitochondria, where both mCherry and GFP signals are observed, resulting in a yellow color. When mitochondria are subjected to mitophagy and fused with lysosomes, only the mCherry signal is detected since the GFP signal is quenched due to the acidic pH of the lysosomes. The mitophagy index was calculated based on the degree of GFP and mCherry co-localization as measured by Pearson coefficient. To determine the effect of PLK1 on mitophagy activation, we transfected the CrT shCtl cells and CrT shPLK1 cells with the mitophagy indicator. An additional group of CrT shCtl cells treated with FCCP were used as a positive control, indicating the level of mitophagy activation. Two days after transfection, we quantified the co-localization of GFP and mCherry. As shown in Figure 4F, mitophagy activation was significantly inhibited in cells with PLK1 downregulation. Meanwhile, we confirmed that knock-down of PLK1 halted the accumulation of poly-ubiquitination on mitochondrial and conjugation of ATG5-ATG12, indicating that knock-down of PLK1 inhibits the activation of mitophagy (Figure S3E). In addition, we testified that knock-down of PDHA1 facilitated the mitochondrial poly-ubiquitination and stabilization of Parkin on mitochondria, further suggesting that PLK1-assoicated PDHA1 downregulation promotes activation of mitophagy (Figure S3F). To further confirm the functional importance of PLK1 on mitophagy, CrT shPLK1 cells were co-transfected with the mitophagy indicator and three different forms of PLK1 constructs (WT, T210D and K82M), respectively. After 2 days, cells were harvested for mitophagy index analysis. As shown in Figure S3G, compared to the cells expressing wild type PLK1, we found no significant difference in mitophagy index in cells expressing PLK1-T210D. We did notice that in some cells with PLK1-T210D, the degree of GFP and mCherry co-localization was dramatically downregulated. Furthermore, mitophagy activation was significantly blocked in cells expressing PLK1-K82M, suggesting that inhibition of PLK1 kinase activity attenuates mitophagy activation in CrT cells. Considering that PDHA1 degradation is closely related to PLK1-mediated phosphorylation and the autophagy/mitophagy-lysosome pathway, we further addressed whether PLK1-associated PDHA1-T57 phosphorylation triggers mitophagy activation. Accordingly, three groups of CrT cells were infected with lentivirus encoding different forms of PDHA1 (WT, T57D and T57A). After puromycin selection, cells stably expressing these PDHA1 variants were transfected with the mitophagy indicator for 2 days, followed by mitophagy analysis. As shown in Figure 4G, mitophagy activation was dramatically promoted in cells expressing phospho-mimetic PDHA1-T57D compared to that of cells expressing wild type PDHA1. As expected, the PLK1 unphospho-mimetic PDHA1-T57A variant had no effect on mitophagy activation. Moreover, mitochondria-lysosome colocalization was significantly upregulated in BEAS-2B cells with PDHA1-T57D expression compared to both cell lines expressing PDHA1 WT or T57A, further implying that phosphorylation of PDHA1 at T57 triggers mitophagy activation (Figure S3H). To further support this, mitochondria were isolated in CrT cells expressing theses PDHA1 variants, and we compared the effects of PDHA1 variants on activation of mitophagy. As shown in Figure 4H, compared to cells expressing wild type PDHA1 and PDHA1-T57A variant, PDHA1-T57D variant promoted mitochondrial poly-ubiquitination, stabilization of P62 as well as PINK1 and conjugation of ATG5-ATG12. In contrast, poly-ubiquitination on mitochondria and stabilization of PINK1 was markedly inhibited in the cells expressing PDHA1-T57A variant. Considering that mitophagy is modulated by multiple regulators, and PDHA1-T57D variant enhances PINK1 and P62 stabilization, we further confirmed whether PLK1/PDHA1-associated mitophagy activation is mediated by PINK1-Parkin-P62 pathway. Thus, Parkin and P62 were knocked down in CrT cells by siRNAs, followed by isolation of mitochondria. As shown in Figure 4I, knocking down of Parkin blocked mitochondrial poly-ubiquitination and further inhibited P62 recruitment on mitochondria, whereas the protein level of PDHA1 was rescued accordingly. Interestingly, although knocking down of P62 also prevented mitochondria from poly-ubiquitination, we noticed that Parkin was stabilized on mitochondria, indicating that PLK1/PDHA1-associated mitophagy is firstly mediated by stabilization of PINK1 and recruitment of Parkin, followed by recruitment of P62. Taken together, PLK1-associated PDHA1-T57 phosphorylation facilitates mitophagy activation, which eventually results in PDHA1 degradation.

### PLK1-mediated PDHA1-T57 phosphorylation affects glucose metabolism and TCA cycle

Glycolysis-derived pyruvate is oxidized to acetyl-CoA by PDH complex and utilized as one of the major carbon sources to maintain TCA cycle. As PLK1 triggers mitophagy activation and PDHA1 degradation in a PDHA1 T57 phosphorylation-dependent manner, we further examined the importance of PLK1-mediated phosphorylation of PDHA1 in glucose metabolism and TCA cycle. We therefore carried out a SIRM analysis of CrT cells expressing PDHA1 variants (T57A and T57D) cultured with uniformly labeled ^13^C ([U-^13^C]-) glucose followed by sample processing and metabolites extraction for nuclear magnetic resonance (NMR) and ion chromatography-fourier transform mass spectrometry (IC-FTMS). ^13^C-enriched metabolites from glycolysis (Figure 5Aa-d), TCA cycle (Figure 5Af-l) and amino acid synthesis (Figure 5Am and n) were identified, and their total amounts were determined. As expected, PDHA1 T57 phosphorylation did not affect glucose uptake (Figure S4A). Analysis of metabolites involved in glycolysis showed that cells with PDHA1 T57D variant fail to utilize ^13^C-labeled glucose to fulfill the carbon influx for TCA cycle as result of accumulation of ^13^C-labeled pyruvate in cell and medium during the production of acetyl-CoA by PDH. In particular, cells expressing PDHA1 T57D variant had low level of glucose 6-phsophate (G6P), fructose 6-phosphate (F6P), phosphoenol-pyruvate (PEP) and lactate (Figure 5Aa, b, c and e) derived from ^13^C-labeled glucose, but a considerably higher level of pyruvate (Figure 5Ad) compared to cells with PDHA1 T57A variant expressed. Taking into account the level of ^13^C-labeled pyruvate was increased both in culture media (Figure S4B) and cell extracts (Figure 5Ad) of the cells expressing PDHA1-T57D variant, confirming the blockage of pyruvate metabolism. In addition to being reduced by NADH to lactate by LDH, pyruvate mainly relies on PDH and PC to enter the TCA cycle and provide carbon for subsequent reaction. We first checked the relative activity of PDH and PC in cells expressing PDHA1 T57A or T57D variants. Citrate m+2/pyruvate m+3 can serve as a surrogate for PDH, while the citrate m+3/pyruvate m+3 ratio can be a surrogate for PC. Our findings support the hypothesis that phosphorylation of PDHA1 at T57 leads to suppressed PDH activity (Figure 5B). To our surprise, PLK1-mediated phosphorylation of PDHA1 halved the catalytic capacity of PC even though the carbon entry into TCA through PC was limited in this case. To assess further whether PDHA1 phosphorylation of T57 affects downstream metabolism of pyruvate, we measured the amounts of several metabolites in TCA cycle. The first turn of the TCA produces metabolites m+2, and this was not affected by the T57D variant expression; subsequent turns producing metabolites m+3, 4 and 5 showed decreases upon T57D variant expression. Most of the significant ^13^C-enriched isotopologues of citrate, cis-aconitate, isocitrate and succinate (Figure 5Af-h and j) were found dramatically decreased in cells with PDHA1-T57D compared to its phospho-null mimics, indicating that phosphorylation at T57 of PDHA1 blocks carbon influx from ^13^C-labeled pyruvate. Particularly, non-glucose substrate contribution, measured through m+0 levels, was considerably higher in cells expressing PDHA1-T57D variant compared to its phospho-null mimics, demonstrating that PLK1-mediated phosphorylation of PDHA1 at T57 prevents cells from utilization of carbon source derived from ^13^C-labeled glucose. By contrast, cells expressing phosphor-null mimics of PDHA1 efficiently consumed ^13^C-enriched pyruvate derived from [U-^13^C]- glucose and led to ^13^C-labeled carbon entering the TCA cycle. Although most of ^13^C-labeled carbon in TCA precursor metabolites originated from [U-^13^C]-glucose such as citrate, cis-aconitate and isocitrate (Figure S4G-I) in cells expressing PDHA-T57A variant, we still observed a shift that nearly half of α-KG, fumarate and malate (Figure S4J, M and N) come from non-^13^C labeled sources, indicating dominant contribution from other carbon sources other than glucose. Similar effects could also be observed in cells expressing PDHA1-T57D variant that more than 40% of TCA precursor metabolites (citrate, cis-aconitate and isocitrate) and 60% of α-KG, succinate, fumarate and malate relied on unlabeled carbon sources. Apart from glucose, glutamate and aspartate can also input carbon into TCA cycle by transamination. To reveal whether glutamate and aspartate can account for unlabeled TCA cycle metabolites, we measured the level of aspartate and glutamate in cells expressing PDHA1-T57A or T57D variants. We found both cell lines had higher level of unlabeled aspartate and glutamate than labeled isotopologues (Figure 5Am and n). More surprisingly, cell expressing PDHA1-T57D variant had a much higher level of both aspartate and glutamate than the cells with phosphor-null mimics, indicating that PLK1-mediated PDHA1 phosphorylation may enhance metabolic reliance on glutamate and aspartate to compensate for the reduced carbon input from pyruvate due to inhibition of PDH function. To better visualize the carbon influx affected by PLK1-mediated PDHA1 phosphorylation, we calculated some of the major ^13^C-enriched metabolic intermediates in glycolysis and TCA cycle according to the carbon metabolic pathways in the cell and displayed the major carbon flows according to their fraction labeled by ^13^C in the cells expressing PDHA1 T57A or T57D variant (Figure 5C). Combining the actual content of these metabolic intermediates, we found that more than 95% of G6P, F6P and PEP (Figure S4C-E) are derived from [U-^13^C]-glucose, but the above metabolites are metabolized and consumed in cells expressing PDHA1 T57D variant at a faster rate than cells expressing PDHA1 T57A (Figure 5Aa-c). For pyruvate, although the ^13^C-enriched pyruvate accounted for 80% of the total content in both cell lines expressing PDHA1 T57A or T57D variants (Figure S4F), the actual content of both labeled and unlabeled pyruvate in cells expressing PDHA1 T57A variant was three times higher than in cell expressing its phosphor-null mimics (Figure 5Ad). Meanwhile, several TCA intermediates in cells expressing PDHA1 T57A variant are derived from [U-^13^C]-glucose, such as citrate, cis-aconitate and isocitrate, the corresponding fraction of ^13^C-labeling in cells expressing the phospho-mimics is only 60% (Figure 5C and Figure S4G-I). This implies that in cells expressing PDHA1 phospho-mimics, the carbon carriers derived from U^13^C-glucose cannot efficiently enter the TCA cycle through the pyruvate metabolic pathway, while cells expressing phosphor-null mimics can use the [U-^13^C]-glucose-derived pyruvate to provide sufficient carbon supplies for the TCA cycle. And this further indicates that PLK1 affects carbon influx by phosphorylating PDHA1 at T57 and ultimately impacts oxidative phosphorylation. In addition, as in our previous report, we observed additional carbon import via glutamate and aspartate in this setting.

**Figure 5.**
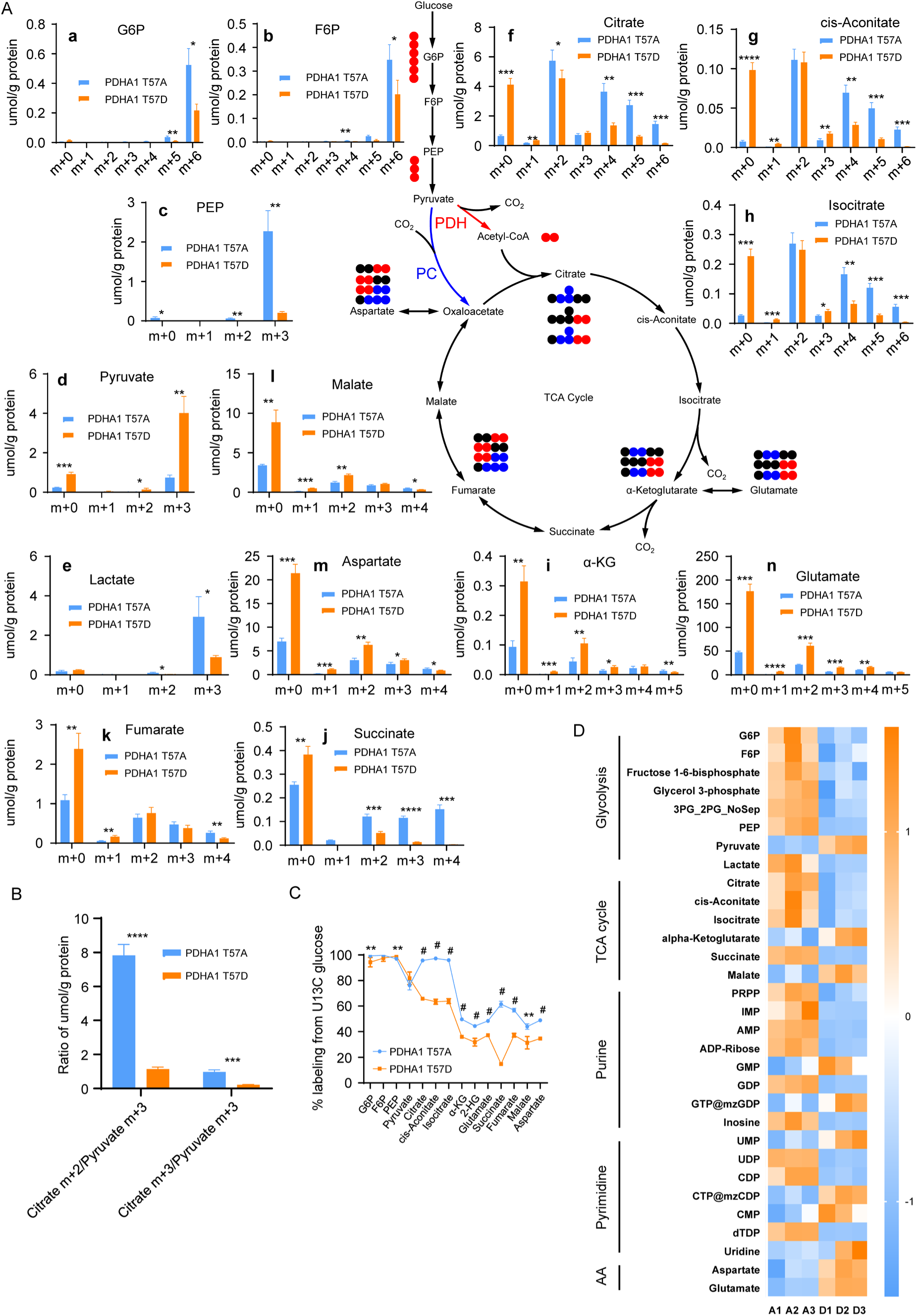
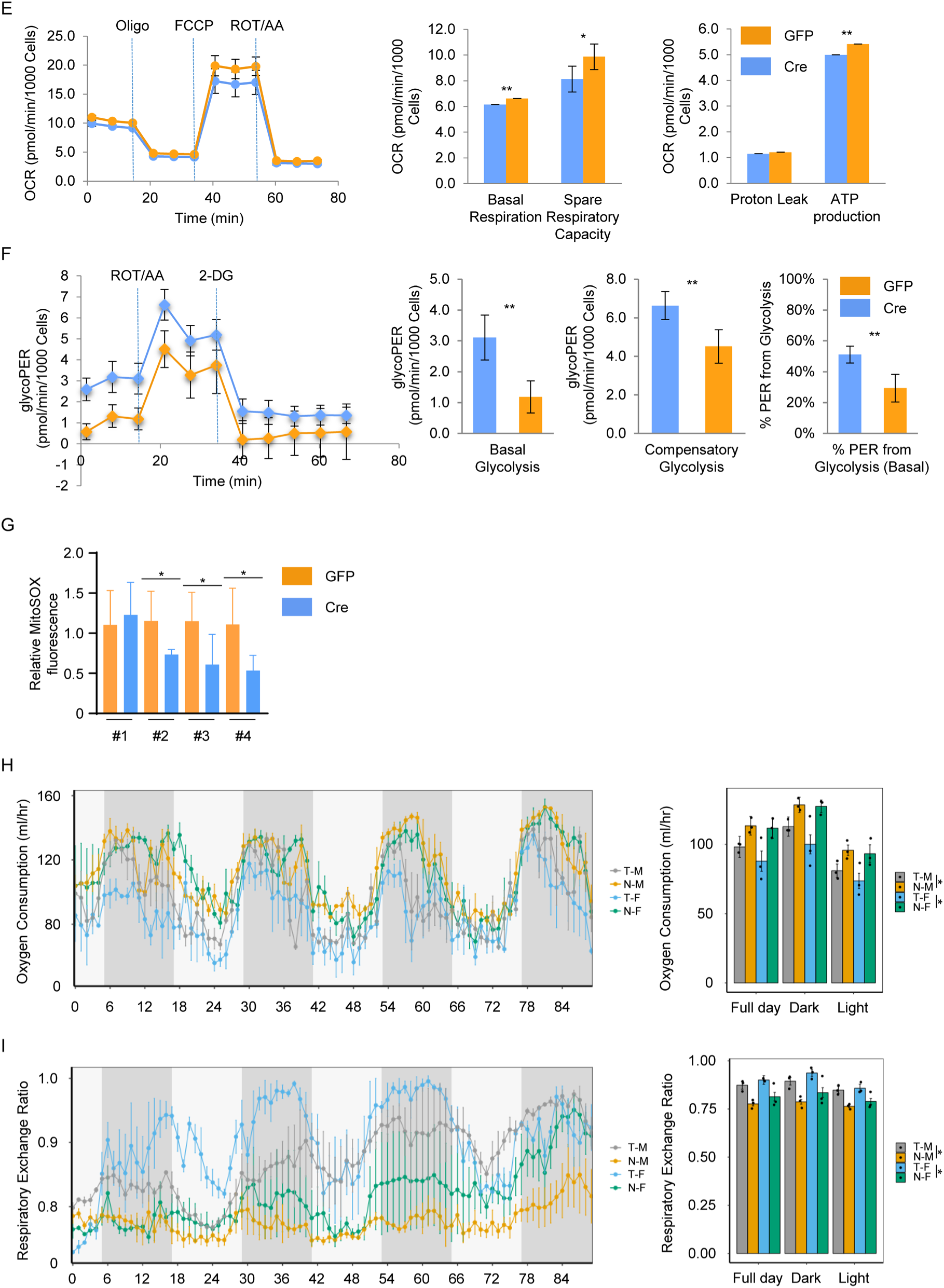
PLK1-mediated PDHA1-T57 phosphorylation affects glucose metabolism and TCA cycle. (A) schematic and quantitative describing the fate of U^13^C-glucose in CrT cells used for glycolysis and TCA-cycle. a-e, f-l and m-n are the total amounts of the isotopologues in metabolites derived from glycolysis, TCA cycle and the aspartate-malate shuttle respectively. Black circle: ^12^C; red circle and blue circle: ^13^C from PDH and PC-derived TCA cycle, respectively. G6P: glucose-6-phosphate; F6P: fructose-6-phosphate; PEP: phosphoenolpyruvic acid; α-KG: alpha-ketoglutarate. The x-axis denotes the number of ^13^C atoms present in each compound. The total amount of isotopologues are presented as means ± standard deviations (n=3), *P<0.05; **P<0.01; ***P<0.001; ****P<0.0001. (B) Citrate m+2/ pyruvate m+3 ratio and citrate m+3/ pyruvate m+3 ratio are presented as surrogates for activity of PDH and PC in cells expressing PDHA1 T57A or T57D variant, respectively. (C) Total fractional enrichments of ^13^C-labeled isotopologues in (A). (D) Heat-map of the total amount of ^13^C-enriched metabolites associated with glycolysis, TCA cycle, purine synthesis, pyrimidine synthesis and amino acid metabolism. (E) Mitochondrial stress test of PDHA1-T57D conditional knock-in MEF. MEFs collected from PDHA1-T57D conditional knock-in mice were transfected by Lenti-GFP-control or Lenti-GFP-Cre for 48h, followed by mitochondrial stress analysis. Quantification of metabolic parameters are presented as means ± standard deviations (n=6). *P<0.05; **P<0.01. (F) Glycolytic rate assay of PDHA1-T57D conditional knock-in MEF. MEFs collected from PDHA1-T57D conditional knock-in mice were transfected by Lenti-GFP-control or Lenti-GFP-Cre for 48h, followed by glycolytic rate assay. Quantification of metabolic parameters are presented as means ± standard deviations (n=6). **P<0.01. (G) (H-I) Indirect calorimetry was performed in Pdha1 T57D knock-in male and female mice. Pdha1 T57D knock-in was activated by tamoxifen injection in both male mice (T-M, n=3) and female mice (T-F, n=3). Mice without tamoxifen treatment were used as group control (N-M and N-F, n=3). Oxygen consumption (H) and respiratory exchange ratio (I) for four groups of mice (T-M, N-M, T-F and N-F) monitored for 84 hours at room temperature. *P<0.05.

To gain a more comprehensive understanding of the potential impact of PLK1 phosphorylation of PDHA1 on cellular metabolism, we integrated the actual content of ^13^C-enriched metabolites in cells expressing PDHA1 T57A or PDHA1 T57D variant related to glycolysis, the TCA cycle, purine synthesis, pyrimidine synthesis and amino acid synthesis (Figure 5D). As we expected, PLK1-mediated phosphorylation of PDHA1 resulted in a reversal of pyruvate metabolism, while upregulation of the aspartate-malate shuttle related metabolites suggested a shift in carbon source from pyruvate to amino acids such as glutamate. To our surprise, we also found that certain purine and pyrimidine metabolites were also affected by PLK1-mediated phosphorylation of PDHA1, such as GMP, GTP, UMP, CMP etc., which also demonstrated the close relationship between PLK1 and cellular metabolism. Taken together, considering that PLK1-meidated PDHA1 phosphorylation triggers PDHA1 protein degradation, cells expressing PDHA1-T57D variant can no longer efficiently consume pyruvate and utilize ^13^C-enriched pyruvate as a carbon source to drive TCA cycle, which results in an accumulation of pyruvate and a shifting dependence from pyruvate to backup carbon sources such as glutamate and aspartate. Hence, PLK1 is tightly involved in glucose metabolism and negatively regulates TCA cycle through phosphorylation of PDHA1 at T57.

To better understand the potential effects of PLK1-mediated PDHA1 T57 phosphorylation on energy metabolism in vivo, a Pdha1 T57D conditional knock-in transgenic mouse line was generated. Firstly, we collected mouse embryonic fibroblasts (MEFs) from the pregnant transgenic mice conditionally expressing PDHA1-T57D variant upon Cre-recombinase induction. To determine whether the phosphorylation of PDHA1 T57 results in metabolism reprogramming in vivo, primary MEFs were transfected with Lenti-GFP and Lenti-GFP-Cre accordingly, followed by mitochondrial stress test and glycolytic rate assay. As expected, MEFs transfected with Lenti-GFP-Cre had a lower level of maximum respiratory capacity and ATP production compared to MEFs transfected with Lenti-GFP (Figure 5E). To compensate the reduction of ATP supplied originated from mitochondria, MEFs with Cre induction shifted their reliance from oxidative phosphorylation to glycolysis (Figure 5F). Noting that PDHA1-T57 phosphorylation blocks the carbon influx from pyruvate to TCA cycle, accumulated pyruvate facilitates the reduction of pyruvate and the regeneration of NAD^+^, which results in a 50% increase of proton efflux rate derived from glycolysis in MEFs with Cre induction.

Mitochondrial superoxide generation were also measured in MEFs transfected with Lenti-GFP or Lenti-GFP-Cre and we found that Cre induction resulted in a downregulation of mitochondrial superoxide level in 3 out of 4 MEF lines (Figure 5G). Then, we performed genetic crosses to incorporate Cre recombinase-estrogen receptor T2 (Cre-ER^T2^) into Pdha1-T57D conditional knock-in mouse model to activate T57D mutant knock-in. We extended our investigation upon mouse whole body metabolism to mouse model with mutated Pdha1, starting with tamoxifen treated mice, which have Pdha1-T57D variant expression to mimic the phosphorylation by PLK1. Here, we measured the effect of Pdha1-T57D on oxygen consumption, carbon dioxide production, respiratory exchange ratio and energy expenditure during light phase and dark phase. Tamoxifen-induced Pdha1-T57D expression repressed oxygen consumption and energy expenditure in male and female mice during both dark and light phases (Figure 5H and S4Q), indicating that tamoxifen-induced mice had a lower metabolic rate compared to the mice without induction. However, contrary to our prediction, the expression of Pdha1-T57D did not affect the production of carbon dioxide. Regardless of the light or dark phase, tamoxifen-induced Pdha1-T57D expression had no significant effect on carbon dioxide production in either male or female mice (Figure S4P). In addition, we observed a significant increase in respiratory exchange ratio in tamoxifen injected mice (Figure 5I), implying that mice with Pdha1-T57D knock-in relied more on carbohydrate than lipid oxidation to produce energy. The greater reliance on carbohydrates did not compensate for the suppression of metabolic rate in mice by PDHA1-T57D expression. This is consistent with our in vitro findings, that is, PLK1-mediated PDHA1 T57 phosphorylation inhibits oxidative phosphorylation and forces cells to produce more ATP for cellular activities through glycolysis. The incomplete metabolism of carbohydrates reduces the energy metabolic efficiency of cells, which is manifested in the lower oxygen consumption and energy expenditure in mice when Pdha1-T57D is introduced into the mice.

### PLK1-mediated PDHA1-T57 phosphorylation dysregulates PDH activity and mitochondrial function to promote tumor growth

As a pivotal linker between glycolysis and OXPHOS, PDHA1 controls carbon influx into the TCA cycle and mitochondrial function by catalyzing pyruvate to acetyl-CoA. We have demonstrated that PLK1 negatively regulates mitochondrial activity and that PLK1-associated PDHA1 phosphorylation results in PDHA1 protein degradation and metabolic reprogramming. We then sought to explore whether and how PLK1 affects mitochondrial function in a PDHA1-dependent manner. Thus, we performed mitochondrial stress tests with CrT cells stably expressing different forms of PDHA1 variants (WT, T57A and T57D). As shown in Figure 6A, both basal respiration and spare respiratory capacity were increased in CrT cells expressing wild type PDHA1 or PDHA1-T57A compared to control CrT cells. However, overexpression of PDHA1-T57D failed to rescue mitochondrial function, suggesting that PLK1-mediated inhibition of OXPHOS is due to PDHA1-T57 phosphorylation. To further investigate the function of PLK1-mediated PDHA1 phosphorylation, we predicted the initial structures of wild-type PDHA1 (PDHA1-WT) and PDHA1-T57-phosphorylation (PDHA1-T57pho) based on the sequences published on UniProt (P08559) by using AlphaFold2(*31*). We further investigated the conformation of the PDHA1-WT and PDHA1-T57pho in explicit aqueous solution using molecular dynamics (MD) simulations (Figure 6B). The final configuration of MD simulation showed that the PDHA1-T57pho (red) presents a conformation different from the PDHA1-WT (blue) in both N-terminal and C-terminal. The difference in structural conformation implies a distinct function between PDHA1-WT and PDHA1-T57pho. The flexibility of PDHA1 can be characterized using the root mean square fluctuation (RMSF) of its Cα atoms. The value of ΔRMSFi between PDHA1-WT and PDHA1-T57pho was calculated as RMSFi (PDHA1-T57pho) – RMSFi (PDHA1-WT), where i refers to the Cα atom on amino acid residue i. As shown in Figure 6C, RES1-25 (N-terminal) and RES293-303 shows the ΔRMSF value <0.5nm, indicating that PDHA1-T57pho has a more rigid conformation compared with PDHA1-WT within these regions. We also found that RES 370-390 (C-terminal) presented the ΔRMSF value >0nm, suggesting that more flexible structures are observed in C-terminal when T57 of PDHA1 gets phosphorylated. In consonance with these findings, compared to secondary structure of PDHA1-WT, both RES1-25 and RES293-303 in PDHA1-T57pho present more ordered structures. Figure S5A shows that T57 phosphorylation of PDHA1 induces conversion of coil structures into helix structures in RES1-25 (a and b) and bend structures in RES293-303 (c and d). In contrast, RES370-390 has more β-sheet structures on PDHA1-WT than the PDHA1-T57pho (Figure S5A e and f). This also explains why ΔRMSF≤-0.5nm for RES1-25 and RES293-303, while ΔRMSF value >0nm for RES370-390. We then examined the phosphorylation-mediated changes of tertiary structures of the PDHA1. Residue-residue contact and mean smallest distance of Cα atoms on PDHA1-wt and PDHA1-T57pho were calculated and analyzed (Figure 6D). The residue-residue contact map of PDHA1-T57pho shows the loss of residue contact within both N-terminal and C-terminal. RES293-300 region also shows an increased residue-residue contact but not as significant as N-terminal and C-terminal. To check the stability of the simulation, radius of gyration (Rg) of protein and solvent accessible surface area (SASA) of protein were investigated. The average Rg of PDHA1 was reduced from 2.230nm to 2.096nm upon T57 phosphorylation (Figure S5B). In addition, SASA% shows that the RES57 presents more exposure area on phosphorylated PDHA1 than the wild type PDHA1 (Figure S5C). Together, these data demonstrate that PLK1-mediated T57 phosphorylation of PDHA1 results in PDHA1 protein structure changes, which possibly leads to distinct enzymatic activity and down-stream consequences.

**Figure 6.**
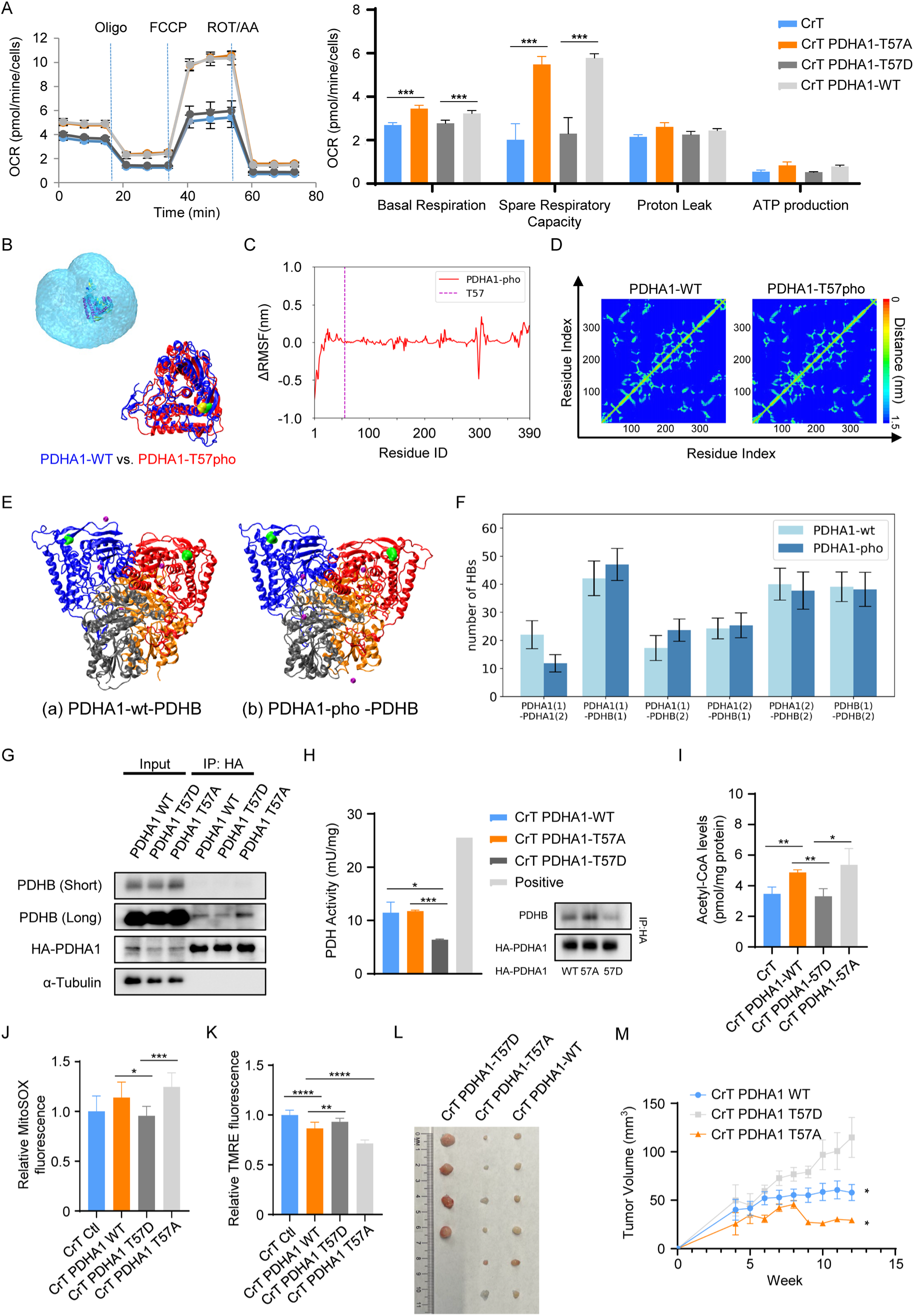
PLK1-mediated PDHA1-T57 phosphorylation dysregulates PDHC activity and mitochondrial function and promotes tumor growth. (A) The mitochondrial stress analysis were assessed in CrT cells stably expressing different forms of PDHA1 constructs (WT, T57D and T57A). Each data point is mean ± standard deviation (n=6); quantification of metabolic parameters are presented as mean ± standard deviations on the right panel. ***P<0.001. (B) The snapshot of a simulation system containing PDHA1-WT and final conformation of PDHA1-wt (blue) and PDHA1-pho (red). The yellow and green surface is the RES57 of PDHA1-WT and PDHA1-T57pho, respectively. (C) ΔRMSF of C_α_ atoms on amino acids of the PDHA1 protein. (D) Mean smallest distance between Cα atoms of amino acid residues on PDHA1-WT and PDHA1-T57pho. (E) The snapshot of the final configuration of (a) PDHA1-wt-PDHB and (b) PDHA1-pho-PDHB heterotetramers. The PDHA1 dimer are bule and red (PDHA1(1)-blue, PDHA1(2)-red), The PDHB dimer are grey and orange (PDHB(1)-grey, PDHB(2)-orange). The K^+^ and Mg^2+^ ions are purple. The RES57 on PDHA1 are green. (F) The average HBs in PDHA1-wt-PDHB and PDHA1-pho-PDHB heterotetramers. (G) Co-IP of HA-PDHA1 WT and HA-PDHA1 variants (T57D and T57A) with PDHB in CrT cells. (H) Quantification of the PDH enzymatic activity in CrT cells expressing HA-PDHA1-WT, HA-PDHA1-T57A and HA-PDHA1-T57D. Samples tested in assay was enriched by IP:HA, followed by IB against HA and PDHB to access the loading volume of samples. (I) Quantification of the acetyl-CoA level in CrT cells stably expressing wild-type PDHA1 and PDHA1 variants (T57D and T57A) are presented as means ± standard deviations (n=3). *P<0.05; **P<0.01. (J) Quantification of mitochondrial ROS level in CrT cells stably expressing wild-type PDHA1 and PDHA1 variants (T57D and T57A) are presented as means ± standard deviations (n=8). *P<0.05; ***P<0.001. (K) Quantification of mitochondrial membrane potential in CrT cells stably expressing wild-type PDHA1 and PDHA1 variants (T57D and T57A) are presented as means ± standard deviations (n=12). **P<0.01; ****P<0.0001. (L and M) Tumor growth of CrT cells stably expressing wild-type PDHA1 and PDHA1 variants (T57D and T57A) in nude mice (n=6). *P<0.05.

The pyruvate dehydrogenase complex (PDHc) is a multi-enzyme complex composed of multiple copies of three enzymes, pyruvate dehydrogenase (E1 component), dihydrolipoamide acetyltransferase (E2 component), and dihydrolipoamide dehydrogenase (E3 component). Considering E1 component is composed of catalytic PDHA1 and regulatory PDHB subunits, and the integrity of this unit is critical for PDH activity, we hypothesized that PLK1-mediated PDHA1-T57 phosphorylation induces a structure change in PDHA1 that might reduce the binding efficiency between PDHA1 and its binding partners. To investigate this, we established the final configuration of WT-PDHA1-PDHB and T57 phospho-PDHA1-PDHB heterotetramer by using AlphaFold2 (Figure 6E) and characterized the interaction of the PDHA1-PDHB complex using protein-protein hydrogen bonds. The hydrogen bonds are defined by the criteria proposed by the Chandler group(*32*). The distance between O (donor) and O (acceptor) is ≤ 0.35 nm. The angle of H (donor)-O (donor)-H (acceptor) ≤ 30°. Table 2 shows the average number of protein-protein hydrogen bonds (NHB) between the four proteins in the complex. The NHB is 22.01±4.99 between two wt-PDHA1 and reduces to 11.82±3.06 between the two pho-PDHA1. No significant reductions are observed between the other unit pairs upon the phosphorylation (Figure 6F and Figure S5D). The nearly 50% reduction of hydrogen bonds between two PDHA1 pairs may cause the decrease of the stability for the complex, further indicating that phosphorylation of PDHA1 at T57 interferes with hydrogen bond formation between two copies of PDHA1 subunit in E1 and resulting in the binding collapse in E1 heterotetramer.

Based on the hints we got from protein simulation, we generated CrT cells stably expressing three HA-tagged PDHA1 constructs (WT, T57D and T57A), which were used to compare the binding capability between PDHB and the PDHA1 variants with co-IP assay. As indicated in Figure 6G, endogenous PDHB was co-IPed with wild type HA-PDHA1. Of note, the phospho-mimetic PDHA1-T57D displayed a lower binding affinity to PDHB compared to that of wild type PDHA1. In contrast, the phosphor-null PDHA1-T57A variant showed a significant increase in binding capability to PDHB. To further determine whether T57 phosphorylation-mediated PDHA1/PDHB binding collapse affects PDH enzymatic activity, PDHA1 protein were enriched from CrT cells stably expressing HA-PDHA1-WT or HA-PDHA1 variant by IP, followed by PDH activity measurement. Compared to PDHA1-WT and PDHA1-T57A, PDHA1-T57D demonstrates decreased PDH activity level, suggesting that PDHA1-T57 phosphorylation blocks PDHA1-PDHB binding and leads to a reduction of PDH enzymatic activity. The protein of PDHA1-WT, PDHA1-T57A and PDHA1-T57D used in PDH enzymatic activity detection were collected, followed by WB. Consistently, we found that PDHA1-T57D variant binds less PDHB compared to PDHA1-WT and PDHA1-T57A (Figure 6H). Noting that PDH catalyzes the conversion of acetyl-CoA from pyruvate, the levels of acetyl-CoA in CrT cells expressing PDHA1-WT and PDHA1 variants were investigated. Cells expressing PDHA1-T57D showed a lower level of acetyl-CoA than that of cells expressing WT and T57A (Figure 6I). MitoSOX accumulates in mitochondria due to its positive charge, reflecting the mitochondrial ROS level. We showed that cells expressing PDHA-T57D had a lower level of mitochondrial ROS than that of cells expressing WT and -T57A (Figure 6J). Tetramethylrhodamine, ethyl ester (TMRE) is a red-orange fluorescent dye that is readily sequestered by active mitochondria due to pre-existing mitochondrial membrane potential. Hyperpolarized mitochondrial membrane potential is chronic feature of cancer, which facilitates cancer cells to antagonize intrinsic apoptosis response and eventually benefits to cancer progression and resistance development of certain anti-cancer drugs(*33*). As indicated in Figure 6K, CrT cells stably expressing PDHA-T57A had the lowest TMRE fluorescence, whereas cells expressing PDHA-T57D had enhanced TMRE fluorescence, indicating that PLK1-mediated PDHA1 T57 phosphorylation promotes hyperpolarization of mitochondria. Consistent results were observed by using HEK293TA cells expressing different forms of tdTomato-PDHA1 constructs (WT, T57D and T57A), followed by TMRE and mitoSOX treatment to indicate mitochondrial membrane potential and mitochondrial ROS generation (Figure S5E and F). These results are not surprising because PLK1-mediated PDHA1 phosphorylation decreases the binding efficacy between PDHA1 and PDHB, which fundamentally impairs PDH activity as well as PDHc-related metabolic flux and eventually undermines mitochondrial physiology. Moreover, these metabolic changes and mitochondrial physiology alterations were largely reversed in cells expressing PDHA1-T57A compared to cells expressing PDHA1-T57D, further demonstrating that PLK1-associated inhibition of mitochondrial function is due to PDHA1 phosphorylation by PLK1.

Having established that PLK1-associated PDHA1 phosphorylation negatively regulates mitochondrial function, we investigated whether PDHA1-T57 phosphorylation has any impact on cancer cell proliferation in vitro and in vivo. We compared the proliferation rate of CrT cells stably expressing different forms of PDHA1 (WT, T57A and T57D). As shown in Figure S5G, expression of PDHA1-T57A led to a marked decrease in cell proliferation compared to cells expressing wild type and PDHA1-T57D. In contrast, expression of PDHA1-T57D promoted cell growth compared to wild type PDHA1. Notably, compared with parental cells without PDHA1 overexpression, cell growth of both cell lines expressing wild type PDHA1 or PDHA1-T57A variant was significantly inhibited, consistent with the notion that PDHA1 promotes mitochondrial ROS generation and enhances mitochondria-associated apoptosis. Finally, in vivo xenograft tumor formation indicated that expression of PDHA1-T57D promoted tumor growth. On the other hand, expression of PDHA1-T57A led to a dramatic reduction of tumor progression (Figures 6L and M and Figure S5H). Taken together, these data suggest that PLK1-mediated PDHA1-T57 phosphorylation impairs mitochondrial function and promotes tumor growth both in vitro and vivo.

### PLK1 inhibition enhances the efficacy of DCA

We reasoned that if rescue of the level of PDHA1 by inhibition of PLK1 is efficient to intensify mitochondrial ROS generation, and subsequently inhibits cancer cell proliferation and cell survival, eventually exacerbate cancer cell apoptosis. It has been reported that DCA is broadly involved in certain cancer therapies by upregulating PDH activity and mitochondria-dependent apoptosis. Therefore, we speculate that activation of PDHA1 enzymatic activity by DCA treatment while upregulating PDHA1 protein level by inhibition of PLK1 might be a novel approach for cancer therapy. To this end, we treated CrT cells (shCtl or shPLK1) with different doses of DCA. As shown in Figure 7A, PLK1 knockdown enhanced the growth inhibition of DCA more than 4-fold in comparison to CrT cells. Using mitoSOX Red, we found that compared with CrT cells, CrT shPLK1 cells showed a significantly higher level of mitochondrial ROS generation upon DCA treatment. However, in CrT shCtl cells, the same dose of DCA had no effect (Figure 7B). The colony formation capacity of CrT shPLK1 cells was significantly inhibited upon DCA treatment compared with CrT shCtl cells (Figure 7C), indicating that downregulation of PLK1 enhances chemo-sensitivity in CrT cells. In addition, inhibition of PLK1 kinase activity increased chemosensitivity to DCA and further induced cell apoptosis in CrT cells, as determined by colony formation and IB (Figures 7D and 7E). Similarly, the level of cleaved-PARP was dramatically increased in A549 cells upon a combination of BI6727 and DCA treatment compared with single treatments (Figure S6A). We also tested most promising predicted combinations for CrT cells in 6*6 dose matrix with DCA and another potent PLK1 inhibitor NMS-P937, followed by the quantification of synergy basing on zero interaction potency (ZIP) synergy score. We observed strong synergy between DCA and NMS-P937 in CrT cells crossing a wide range of concentrations (Figure 7F and G). We showed that the maximum ZIP synergy score exceeds 22, which indicates a 22% additional inhibition due to the synergy compared to mono-treatment of two drugs. Next, we treated nude mice carrying CrT xenograft tumors with the PLK1 inhibitor NMS-P937, DCA or both for 3 weeks.

**Figure 7.**
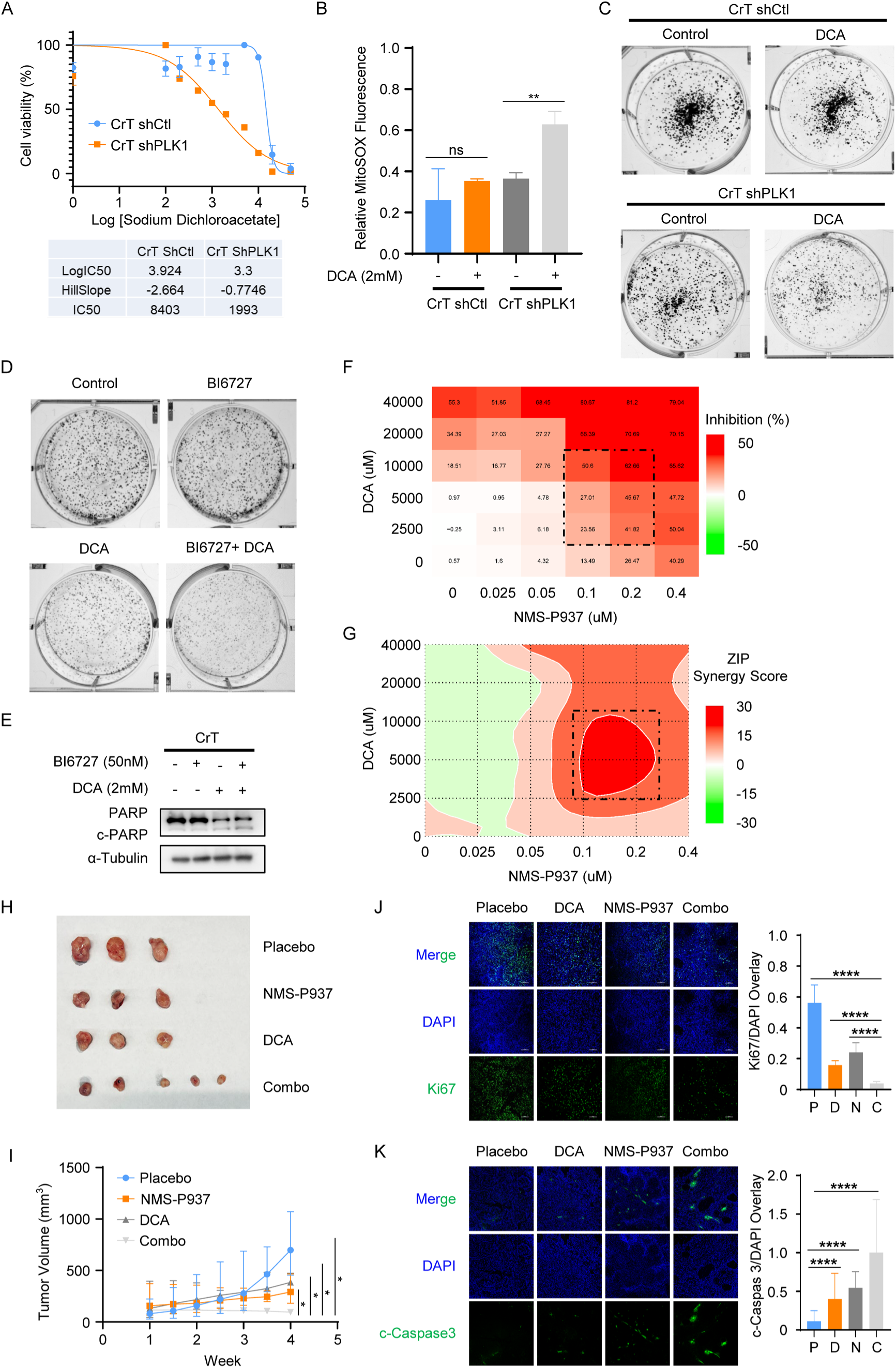
PLK1 inhibition enhances chemosensitivity of DCA in vitro and in vivo. (A) MTT analysis of CrT cells (shCtl or shPLK1) that were treated with increasing doses of DCA is presented as means ± standard deviations (n=3). (B) Quantification of mitochondrial ROS level in CrT cells (shCtl or shPLK1) that were treated with 2mM DCA is presented as means ± standard deviations (n=3). **P<0.01; ns P>0.05. (C) Representative images of colony formation analysis of CrT cells (shCtl or shPLK1) after 2mM DCA treatment. (D) Representative images of colony formation analysis of CrT cells after 2mM DCA and 2nM BI6727 treatment. (E) CrT cells were treated with 50nM BI6727 and 2mM DCA as indicated, followed by IB analysis of c-PARP protein level. (F) Dose response map showing the inhibition rate of cell viability for CrT cells treated with DCA + NMS-P937. (G) Synergy matrix plot showing ZIP score for CrT cells treated with DCA + NMS-P937 (average ZIP synergy score = 9.58; max ZIP synergy score = 22.7). (H) DCA plus NMS-P937 inhibits tumor growth synergistically. Nude mice were inoculated with CrT cells, randomized into four groups (n=5 per group), and treated with either placebo, DCA, NMS-P937 or both for 4 weeks. Representative images of xenograft tumors upon harvest. (I) Tumor volumes over the four-week period. *P<0.05. (J) IF staining for Ki67 shows fewer proliferating cells in tumors with DCA and NMS-P937 combination treatment. Quantification of Ki67 staining is presented on the right. ****P<0.0001. (K) IF staining for cleaved-caspase 3, a marker for cell apoptosis, shows more apoptotic cells in tumors with DCA and NMS-P937 combination treatment. Quantification of cleaved-caspase 3 staining is presented on the right. ****P<0.0001.

As shown in Figure 7H and I, NMS-P937 or DCA alone slightly decreased tumor volume, whereas the combination of both drugs dramatically decreased tumor volume. This regiment had no effect on body weight (Figure S6B). As shown in the representative images of H&E staining, tumors from mice treated with a combination of DCA and NMS-P937 revealed decreased cellular density and increased distribution of apoptotic bodies compared to single treatments (Figure S6C). We also demonstrated that the combination of DCA and NMS-P937 halted tumor cell proliferation and induced a higher level of cell apoptosis as indicated by IF staining against Ki67 and cleaved-caspase 3, suggesting that NMS-P937 plus DCA effective by inhibiting Cr (VI)-associated cancer progression, and that inhibition of PLK1 enhances the efficacy of DCA in vivo (Figures 7J and 7K).

### High PLK1 and low PDHA1 is predictive of poor clinical outcome in NSCLC

Aberrant upregulated PLK1 in cancer cells influences oxidative phosphorylation by promoting PDHA1 degradation, resulting in disorder of mitochondria and increased cancer progression. We then investigated whether such PDHA1 dependent mechanism for metabolic reprogramming by PLK1 would apply to clinical samples. Considering that Cr(VI)-associated lung cancers are squamous cell carcinomas of the central airway(*34, 35*) and adenocarcinomas of the peripheral airways(*36*), we performed immunohistochemistry staining with anti-PLK1 and anti-PDHA1 antibodies in a tissue micro array (TMA) containing 216 patient samples of NSCLC from a clinical cohort. As shown in Figure 8A, we observed a negative correlation between the protein level of PLK1 and PDHA1, supporting the idea that PLK1 mediates PDHA1 downregulation both in vitro and in patients. Furthermore, a higher level of PLK1 and a lower level of PDHA1 correlated with poorly differentiated tumors in patients (Figure 8B), indicating that the PLK1/PDHA1 axis is indeed involved in tumor differentiation and aggressiveness. This was also supported by the Kaplan-Meier analysis (Figure 8C), and survival curves were generated to indicate the preliminary conclusions regarding the influence of the PLK1/PDHA1 axis on the prognosis of patients with NSCLC. A high expression of PLK1 was significantly associated with a worse overall survival for NSCLC cases. In addition, a low expression of PDHA1 was associated with a worse overall survival in the dataset. Taken together, these findings support the rationale to test the PLK1/PDHA1 axis as diagnostic targets to predict the clinical outcome in NSCLC patients.

**Figure 8.**
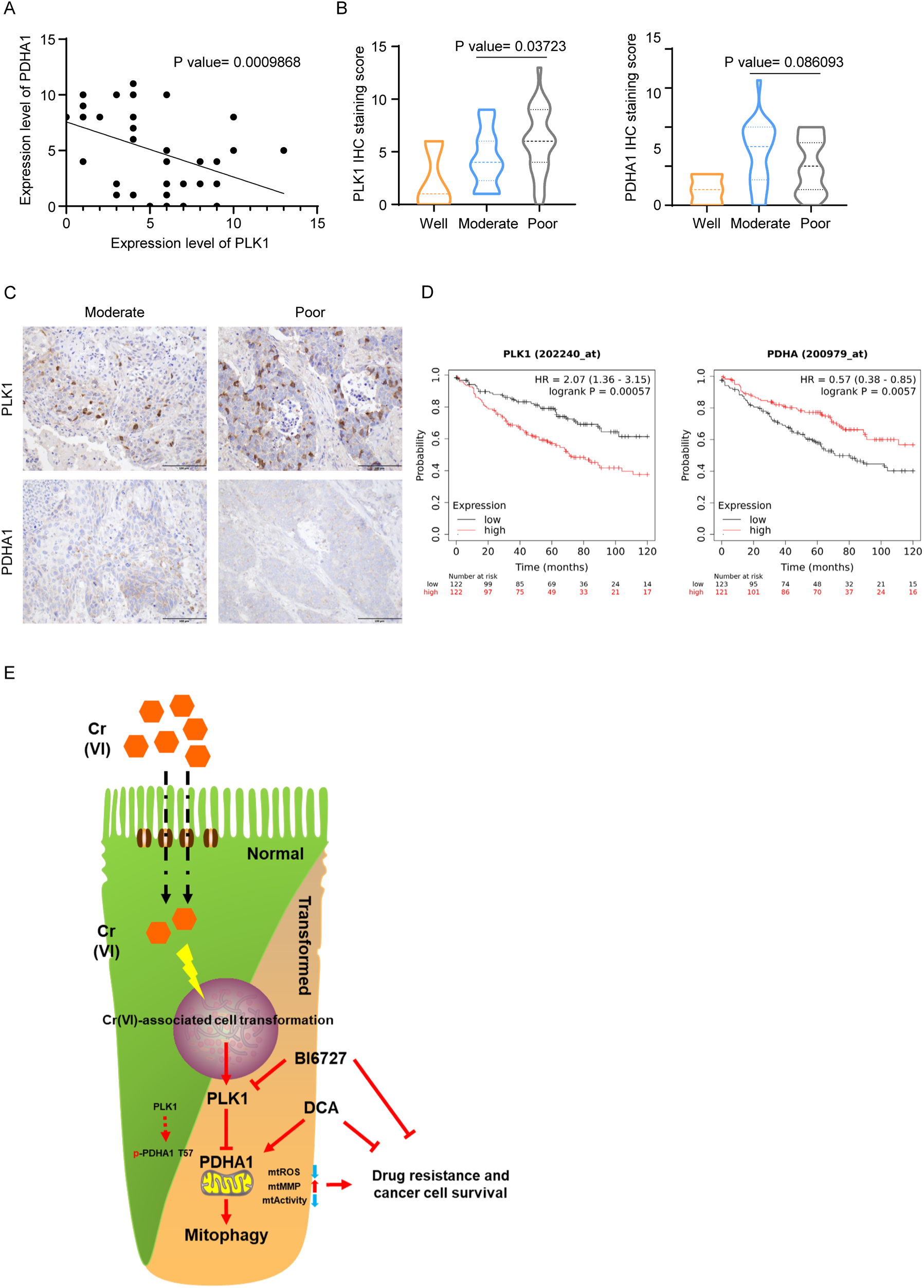
High PLK1 and low PDHA1 is predictive of poor clinical outcome in NSCLC. (A) The protein expression levels of PDHA1 and PLK1 were scored and evaluated, and the correlation between the two proteins is shown. (B and C) IHC staining of PLK1 and PDHA1 was performed in a TMA containing 216 patient samples of NSCLC from a clinical cohort. PDHA1 expression results in granular cytoplasmic staining whereas PLK1 expression results in nuclear and cytoplasmic staining. The reciprocal staining pattern between PDHA1 and PLK1 is shown in representative moderate and poorly differentiated samples. The protein expression levels in NSCLC with well, moderate, and poor differentiation are shown. Note the statistically significant (*P<0.05) increase PLK1 staining as differentiation decreases. (D) Kaplan-Meier-10-year overall survival of NSCLC patients was analyzed according to the expression level of PLK1 and PDHA1. Note the decreased probability of survival with high PLK1 expression (***P<0. 001) and low PDHA1 expression (**P<0. 01). (E) PLK1 inhibition enhances chemo-sensitivity of DCA in Cr(VI)-associated cancer progression.

## Discussion

During cancer progression, cancer cell metabolism undergoes dynamic adaptation to meet the energy demands of cancer cells for proliferation, invasion, metastasis, and antagonizing the activation of apoptotic pathways activated by environmental stresses. Metabolic reprogramming, a common feature of cancer cells, is also prominently present in Cr(VI)-associated cell transformation and Cr(VI)-associated cancer development. In Cr(VI)-associated cell transformation, there is a significant increase in cellular glycolysis, along with inhibition of mitochondrial function and reduction of OXPHOS, which further promote cancer progression. However, most of the research on Cr(VI)-associated cell transformation has focused on the effects of short-term or chronic Cr(VI) treatment on glycolysis, with limited investigation into the inhibitory mechanisms on mitochondrial function and OXPHOS. In our study, we identified polo-like kinase 1 (PLK1) as a key player in Cr(VI)-mediated cell transformation and cancer progression, and as a link between Cr(VI)-associated cancer progression and OXPHOS reprogramming. PLK1 is a protein kinase known for its role in maintaining cell cycle homeostasis, and it has also been implicated in other physiological events, including cell metabolism. PLK1-mediated metabolic reprogramming has been consistently demonstrated in multiple cancer systems including prostate cancer, melanoma, and colon cancer etc(*21, 27*). However, the mechanisms underlying PLK1’s involvement in OXPHOS regulation are poorly understood. In our study, we used lentivirus-based approaches or treatment with a PLK1 inhibitor to show that inhibition of PLK1 results in upregulation of OXPHOS in both CrT and H2009 cells, indicating that this observation is not limited to a specific cell line. Furthermore, overexpression of PLK1 in BEAS-2B cells led to inhibition of mitochondrial activity and contributed to tumor formation and growth, supporting the role of PLK1 in modulating mitochondrial function in the context of Cr(VI)-associated cancer progression.

To dissect the underlying mechanism of PLK1-associated inhibition of OXPHOS, we identified PDHA1 as the major downstream target in this process. As one of the most critical catalytic enzymes functioning in the metabolic flow from glycolysis to OXPHOS, PDHA1 has a well-documented role on cancer metabolism. Accumulating evidence has shown that the enzymatic activity and protein level of PDHA1 are significantly downregulated in a variety of cancers, including ovarian cancer, gastric cancer, etc(*37, 38*). Furthermore, it has been shown that normalization of glucose metabolism by stimulating PDH in cancer cells restores their susceptibility to anoikis and impairs their metastatic potential in breast cancer(*39*). We propose a detailed model to explain how upregulation of PLK1 during Cr(VI)-associated transformation induces PDHA1 instability and undermines mitochondrial activities. PLK1 phosphorylates of PDHA1 at T57, subsequently promoting PDHA1 protein degradation. Using site-specific phospho-antibodies, we demonstrated that PDHA1-T57 phosphorylation indeed occurs in vivo in a PLK1-dependent manner.

Under normal physiological conditions, cells tend to maintain a intact mitochondrial network to efficiently supply energy. However, in cancer cells, mitochondrial dynamics are altered to adapt to increased energy demands and environmental stress. Elevated levels of PLK1, consistent with its oncogenic role, have been shown to induce mitochondrial fission and compromise mitochondrial function, likely through phosphorylation of PDHA1. Mitochondrial fission is considered a precursor to mitophagy, a selective form of autophagy where damaged mitochondria are targeted for degradation in autophagolysosomes. Mitochondrial fission is considered as a pre-step for mitophagy(*40*) that is frequently reported to drive malignant phenotypes of cancer. Mitophagy is a selective mode of autophagy in which mitochondria are specifically targeted for degradation at the autophagolysosome, which helps the recycle of damaged mitochondria or dysfunctional proteins and enhances cancer progression. Herein, we noticed that PDHA1-T57 phosphorylation resulted in PDHA1 translocation from mitochondria to cytosol, likely due to phosphorylation-induced binding collapse between PDHA1 and PDHB in E1 and mitochondrial dysfunction mediated activation of mitophagy. Furthermore, we determined that PDHA1-T57 phosphorylation-mediated degradation is accomplished mainly by activation of mitophagy, as PDHA1 levels were restored by treatment with several autophagy/mitophagy inhibitors.

The study also revealed the potential link between the cell cycle and cell metabolism via PLK1-PDHA1 axis. Our findings in CrT cells expressing PDHA1 T57A variant and PDHA1 T57D variant strongly support our hypothesis that PLK1-mediated PDHA1 phosphorylation triggers metabolic switch from oxidative phosphorylation to glycolysis in vitro. It is also notable that cells with PDHA1 phospho-on mimics rely more on aspartate-malate shuttle, rather than glucose-derived pyruvate to maintain the TCA cycle. Moreover, the metabolic shifting was also found in both MEFs and transgenic mice conditionally expressing PDHA1-T57D variant, highlighting the effect of PLK1 on metabolism reprogramming in vivo. While we did not observe a significant change in oxidative phosphorylation upon Cre induction, we did observe a dramatic promotion of glycolysis. In MEF cells, nearly 70% of the energy is supplied by mitochondria (Figure 6K), and pyruvate is not the only source of carbon input for TCA cycle. Hence, inhibition of pyruvate metabolism via phosphorylation of PDHA1 by PLK1 maybe not sufficient to completely shut down oxidative phosphorylation. Similar observations were made in whole-body metabolism tests and calorimetry experiments in mice using isotopic tracers, supporting the compensatory role of the aspartate-malate shuttle in the presence of PLK1-mediated PDHA1 T57 phosphorylation, which leads to accumulation of pyruvate and increased lactate production, tipping the energy balance towards glycolysis. Notably, these metabolic changes are accompanied by alterations in mitochondrial function and cell cycle-related activities, as previous studies have shown that mitochondrial function and morphology are dynamically regulated during the cell cycle. Furthermore, this regulatory relationship is closely linked to cancer development in cancer cells, as PLK1 not only plays a key role in cell cycle regulation but is also frequently implicated in cancer development. Elucidating the function of PLK1 in oxidative phosphorylation in cancer cells is expected to enhance our understanding of the dynamic interplay between cell cycle and cell metabolism.

Considering the inhibitory effect of PDHA1 on cancer progression, we speculate that it is feasible to simultaneously target PLK1-dependent PDHA1 degradation and PDHA1 activity for cancer therapy. Previous studies have shown that dichloroacetate (DCA), a PDHA1 activator, is involved in certain cancer therapies by upregulating PDH activity and promoting mitochondria-dependent apoptosis. Even with attempts to integrate DCA into cancer treatment, DCA has not been incorporated into clinical cancer treatment regimens due to excessively high effective doses and possible peripheral neuropathy(*41*). Given that PLK1 mediates PDHA1 protein degradation and mitochondrial dysfunction, we speculate that inhibition of PLK1 could enhance the efficacy of DCA in cancer therapy by upregulating PDHA1 protein levels (Figure 8E). Moreover, our results demonstrate that cancer cells with downregulated PLK1 are four-fold more sensitive to DCA treatment, supporting the rationale for a novel combination therapy using a PDHA1 activator like DCA and a PLK1 inhibitor. Our in vivo data from xenograft models also support this notion. These findings suggest that a future clinical trial combining a PLK1 inhibitor and a PDHA1 activator could be a promising strategy for cancer therapy.

## Materials and Methods

### Cell culture, chemicals and reagents

Human bronchial epithelial cell line BEAS-2B, human embryonic kidney 293TA cell line and lung cancer cell lines H2009, H2030 and A549 were obtained from the American Type Culture Collection (Rockville, MD). Chromium-transformed (CrT) cells were generated as described (*42*). BEAS-2B, HEK-293TA, CrT and A549 cells were cultured in Dulbecco’s modified Eagle’s medium (DMEM) supplemented with 10% fetal bovine serum (FBS) and 1% penicillin/streptomycin at 37°C in a humidified atmosphere with 5% CO_2_ in air. H2009 and H2030 cells were cultured in RPMI 1640 supplemented with 10% fetal bovine serum (FBS) and 1% penicillin/streptomycin at 37°C in a humidified atmosphere with 5% CO_2_ in air. Sodium dichromate dehydrate (Thermo Fisher Scientific) was used for Cr(VI) treatments. For short-term exposure to Cr(VI), cells were grown to 80–90% confluent, and then the medium was replaced with DMEM medium containing sodium dichromate dehydrate at the dose and duration indicated. BI6727, MG132, cycloheximide, DCA, NMS-P937, 3-MA, chloroquine, liensinine and doxycycline were obtained from Selleckchem.

### Antibodies

Antibodies against PLK1, PDH, Flag, phospho-Histone H3 (Ser10), PDHB, PARP, cleaved-Caspase 3, VDAC, GAPDH, β-actin and α-tubulin were purchased from Cell Signaling Technology (Danvers, MA) and antibodies against tdTomato and Ki-67 were obtained from Origene (Rockville, MD) and Abcam (Boston, MA), respectively. Antibodies against phospho-PDHA1 (T57) were generated by SinoBiological (Wayne, PA). In brief, a peptide containing phosphorylated-T57 was used to immunize two rabbits over a two-month period. Serum was collected, followed by affinity chromatography-based purification to obtain pT57-PDHA1 antibodies.

### Immunoblotting

Upon harvest, cells were suspended with TBSN buffer (20 mM Tris-HCl, pH 8.0, 0.5% NP-40, 5 mM EGTA, 1.5 mM EDTA, 0.5 mM sodium vanadate, and 150 mM NaCl) with protease inhibitors and phosphatase inhibitors, sonicated and then centrifuged, and protein concentration was measured by Protein Assay Dye Reagent from Bio-Rad. Equal amounts of protein from each sample were mixed with SDS loading buffer, resolved by SDS-PAGE, and transferred to PVDF membranes, followed by incubations with appropriate primary and secondary antibodies.

### Cell viability assay

Cells were seeded onto 96-well plates and incubated overnight, then treated with the respective agents for an additional 3 days. Cell viability was determined using AquaBluer (MultiTarget Pharmaceuticals LLC) as an indicator of viable cells according to manufacturer’s recommendation. Each assay consisted of 6 replicate wells and was repeated at least three times. Data are expressed as the percentage survival of the control, which was calculated using fluorescence after correcting for background noise.

### Cell invasion assays

The cell invasion assays were performed using Transwell (6.5-mm diameter, 8-μm pore polycarbonate membrane), which was obtained from Corning (Cambridge, MA). Cells (10^5^) in 200μL medium were placed in the upper chamber, and the lower chamber was filled with 1 mL serum-free medium supplemented with 0.1% bovine serum albumin. After incubation for 24 h, non-migrating cells were removed using cotton swabs, and the cells that migrated to the lower surface of the filter were stained with crystal violet. Cells were counted using a microscope. Triplicate results are expressed as the mean (standard deviation).

### CrT cell-Derived Mouse Xenograft Model

All animal experiments described in this study were approved by the University of Kentucky Animal Care and Use Committee. CrT cells were inoculated subcutaneously into nude mice (Jackson Laboratories). Two weeks later, animals were randomized into treatment and control groups. NMS-P937 and DCA were orally delivered into the stomach twice a week at a dose of 45mg kg−1 and 100mg kg^-1^, respectively. Tumor volumes were calculated from the formula V=L×W^2^/2 (where V is volume [cubic millimeters], L is length [millimeters], and W is width [millimeters]).

### Mitochondrial stress test

Mitochondrial stress test was performed using a Seahorse Bioscience XF96 Extracellular Flux Analyzer (Seahorse Bioscience). 10^4^ cells were seeded in Seahorse XF96 V3 PS cell culture microplates and grown overnight. Before the experiment, cells were washed and changed to seahorse assay medium (supplemented with 10 mM glucose, 1 mM pyruvate, 2 mM glutamine, adjusted to pH 7.4), then incubated in a non-CO_2_ incubator for 1h. After measuring the basal OCR level, oligomycin, FCCP and antimycin/rotenone were sequentially injected into the cell chamber according to Seahorse standard protocol. The basal OCR level was calculated based on the AUC (area under the curve) before the injection of oligomycin. The maximal OCR level was calculated based on the AUC between the injection of FCCP and antimycin/rotenone.

### Quantitative reverse transcription PCR

Total RNA was extracted with the RNeasy Mini Kit (QIAGEN) and treated with DNase (Ambion, Thermo Fisher Scientific). Quantitative reverse transcription PCR (qRT-PCR) mixtures were assembled with 1μl cDNA template, iQ SYBR Green Supermix (Bio-Rad), and primers for PDHA1 and mtCYB. PCR was carried out for 40 cycles with the following thermal cycling conditions: 95°C for 10 seconds (denaturation) and 61°C for 60 seconds (annealing). All data were normalized to β-Actin mRNA expression levels.

### Acetyl-CoA Level Determination

The level of acetyl-CoA was determined by use of a PicoProbe^TM^ Acetyl-CoA Fluorometric Assay Kit (Biovision)(*43*). Briefly, 10^6^ cells were lysed by the corresponding assay buffer. After samples were deproteinized with the 10 kDa molecular weight cut off spin columns (Biovision), specific reactions were performed according to manufacturer’s instructions to measure the fluorescence using a spectrofluorometer.

### In vitro PLK1 Kinase Assay

In vitro kinase assays were performed with TBMD buffer (50 mM Tris [pH 7.5], 10 mM MgCl_2_, 5 mM dithiothreitol, 2 mM EGTA, 0.5 mM sodium vanadate, and 20 mM p-nitrophenyl phosphate) supplemented with 125μM ATP and 10μCi of [γ-^32^P]ATP at 30°C for 30 min in the presence of purified GST-PDHA1 and PLK1. After the reaction, mixtures were resolved by SDS-PAGE, the gels were stained with Coomassie brilliant blue, dried, and subjected to autoradiography.

### Mitochondrial imaging and MINA Network Analysis

BEAS-2B, CrT, CrT shCtl and CrT shPLK1 cells were grown on glass coverslips in a 6-well plate in DMEM medium until 70% confluent. Cells were then washed twice with phosphate-buffered saline (PBS) and pre-incubated with 5 Mitotracker Red (Thermo Fisher Scientific) in PBS for 15 min at 37°C in the dark. After treatment, the dye was removed, and the cells were washed three times with PBS and fixed with ice-cold methanol. Coverslips were then mounted onto glass slides using Antifade mounting medium. Images were taken using a Nikon A1+ confocal microscope. The images generated were preprocessed on ImageJ (NIH, Bethesda, MD, USA) following steps previously outlined (*44*). After pre-processing, the images were skeletonized. Post skeletonization, images were segmented using Adobe Photoshop CC 2018. These segmented images were opened in ImageJ, and the MiNA macros were used to quantify mitochondrial morphological parameters of each segmented image.

### In Vitro Mitophagy Assay

To examine mitophagy, we transfected cells with Su9-mCherry-GFP, and fixed the cells with paraformaldehyde to visualize mCherry and GFP signals. The mitophagy activation index was calculated based on the degree of GFP and mCherry co-localization as measured by Pearson coefficient. Decreased co-localization (i.e. a decrease in mitophagy index) indicates increased mitophagy. Images were taken with a Nikon A1+ confocal microscope and the Pearson coefficient was determined using Nikon NIS-elements software.

### Indirect calorimetry

Oxygen consumption, carbon dioxide production, respiratory exchange ratio and energy expenditure were measured with Sable Promethion (Sable Systems International, North Las Vegas, NV). Indirect calorimetry measurements were recorded for each mouse (n=12) for at least 84 hours, which allows the analysis cover the pre-acclimation and post-acclimation periods. Data generated were analyzed with CalR as described by Mina etal(*45*).

### Molecular model and simulation detail

All-atom models were used to describe the wild-type and phosphorylated PDHA1 protein (PDHA1-wt and PDHA1-pho), water molecules and ions. The initial structures of wild-type PDHA1 and PDHA1-pho were predicted by AlphaFold2(*31*) using the sequences published on UniProt (P08559). The simulation systems are created by placing a PDHA1 protein or variant in the center of a cubic box 24 × 24 × 24 *nm*^3^ and surrounding the protein with a 3.5nm-thick shell of 0.15M NaCl solution. Counter-ions were added to neutralize the system. The simulation box size is determined to ensure that the protein does not interact with its mirror. The 3.5 nm shell of NaCl solution is used to ensure the protein is surrounded by enough aqueous solution. This work deploys the GROMOS 54B7 force field(*46*) to describe bonded and non-bonded interactions in the systems. The non-bonded interactions are a sum of short-range Lennard-Jones 12-6 potential and long-range coulombic potential, as shown in Equation 1. The bonded interactions are a sum of the bond length, bond angle, and dihedral potentials, as described in the force field.

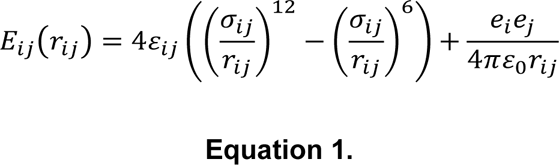

where *E_ij_* is the potential energy due to the non-bonded interactions between atoms *i* and *j*, *r_ij_* is the distance between atoms *i* and *j*, *ɛ*_*ij*_ is the energetic parameter, *σ*_*ij*_ is the geometric parameter and *e*_*i*_ is the partial charge of atom i. The Jorgensen mixing rule is applied to obtain *ɛ*_*ij*_ and *σ*_*ij*_ for atoms belonging to different types.

**Table. 1.** Details of the two systems of PDHA1 in NaCl solution.

| System | Protein | Number of water molecules |
| --- | --- | --- |
| PDHA1-wt | wild-type PDHA1 | 45377 |
| PDHA1-PHO | phosphorylated PDHA1 | 45540 |

**Table 2.**
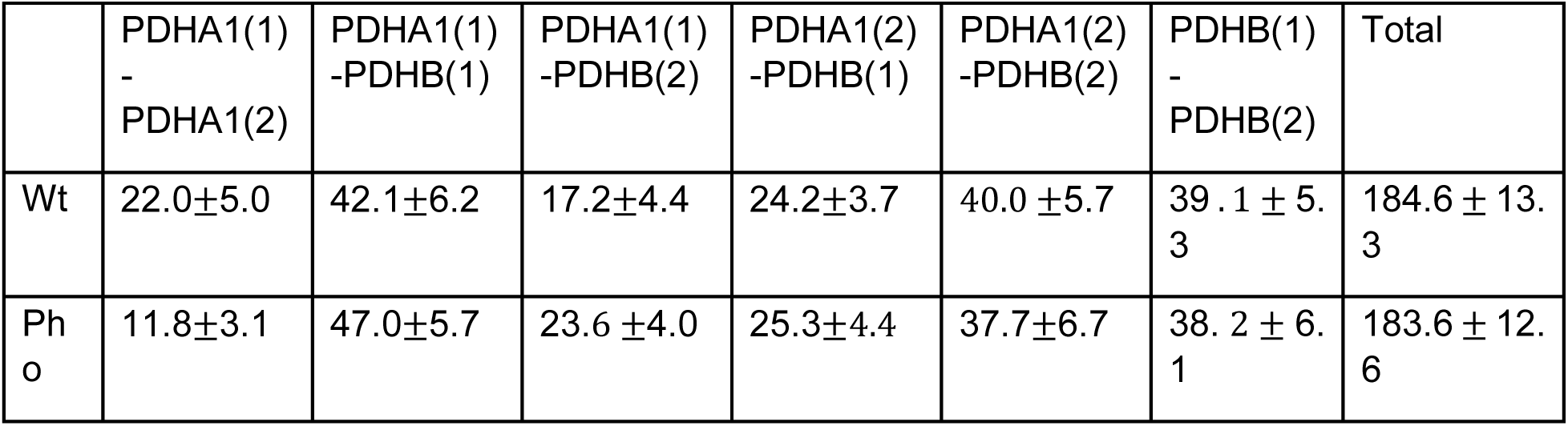
The average number of HBs in PDHA1-wt-PDHB and PDHA1-pho-PDHB heterodimers.

A three-step simulation process was conducted for every simulation system. First, energy minimization was conducted to remove any too-close contacts between atoms. Second, a 200-ns canonical (NVT, T=310 K) ensemble MD simulation (integral step = 2 fs) was conducted to let the system reach thermodynamic equilibrium. Third, a 300-ns canonical (NVT, T=300 K) ensemble MD simulation (integral step = 2 fs) was conducted to collect the trajectory at a frequency of 200 ps. The Berendsen method(*47*) is used to control the temperature and pressure of the system in the second step because it allows the system to reach the desired pressure and temperature at a fast pace. The velocity-rescaling method(*48*) is used to control the temperature of the system in the third step. The short-range van der Waals interactions use a 1.2-nm cut-off, and the long-range electrostatic interactions were calculated using the particle mesh Ewald sum.(*49*) All bonds involving H atoms were constrained during the simulations. The energy minimization and MD simulations for all the systems were conducted using Gromacs-2021.(*50*)

### Mitochondrial-ROS detection

Cells were seeded and grown in a 96-well plate in DMEM medium until 70% confluent. Cells were then washed twice with PBS and pre-incubated with 5 μM MitoSOX Red (Thermo Fisher Scientific) in PBS for 15 min at 37°C in the dark. MitoSOX Red allows selective visualization of O_2_•− generated in the mitochondria because it is rapidly oxidized by O_2_•− only. After treatment, the dye was removed, and the cells were washed three times with PBS. Fluorescence was measured at 530/590 nm by a TECAN plate reader.

### Colony formation assay

Cells (10^3^) were seeded in 6-well plates with 2 mL DMEM supplemented with 10% FBS. One day later, the medium was replaced with new medium containing different drugs. After 6 days, the colonies were fixed by 10% formalin and stained with 5% crystal violet. Colony numbers were counted by using ImageJ software.

### Immunofluorescence (IF) analysis

Xenograft tumors were subjected to paraffin embedding after fixation with 10% formaldehyde overnight. After dewaxing, the sections were heated at 95°C in pH 6.0 citric acid solution for epitope retrieval, followed by permeabilization with 0.2% Triton X-100 and quenched in 3% H_2_O_2_ to abolish endogenous peroxidase. After eliminating peroxidase, tumor tissues were blocked with 10% horse serum in PBS, and incubated with antibodies against Ki67 (1:200), and cleaved-caspase 3 (1:200) in buffer A (1% BSA, 0.1% Triton X-100, 10% horse serum in PBS) for 30 min at 37°C. Tumor tissues were then incubated with Alexa Fluor 488 goat anti-rabbit secondary antibody and mounted using DAPI. The cells were visualized using a Nikon A1+ confocal microscope.

### Stable Isotope Tracer Metabolic Analyses

Cells were grown in DMEM medium containing 25mM [U-13C]-glucose on 10 cm plates in triplicate. Media samples (50 μL) were taken at 3, 6, 12, 24h and immediately flash frozen in liquid nitrogen. Cells were quenched using CH_3_CN and extracted using a modified Folch method as previously described(*51*). Chloroform was added into the mixture of CH3CN and water. Polar fraction, lipids fraction and protein were enriched and extracted from CH3CN-chloroform mixture for stable isotope tracer metabolic analyses.

### NMR

Polar extracts were reconstituted in 35 μL D_2_O containing 17.5 nmol DSS as internal chemical shift reference and concentration standard. ^1^H PRESAT and ^1^H{^13^C}-HSQC spectra were recorded at 15 °C on a 16.45 T Bruker Avance III spectrometer equipped with a 1.7 mm inverse triple resonance cryoprobe as previously described(*52*). Raw data were processed and analyzed using MRestNova. Metabolites and isotopomers were identified using in-house databases (*53*), and quantified as μmol/mg protein and as fractional enrichments F as:

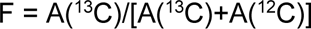

Where A(^13^C) is the peak area of a proton attached to ^13^C, and A(^12^C) is the peak area of a proton attached to ^12^C.

### Uhr IC-FT-MS Analysis

Ion chromatography-ultra high-resolution-Fourier transform-MS (IC-UHR-FTMS) was performed as previously described (*54*). Briefly, polar extracts were reconstituted in 20 μL Nanopure water, and analyzed by a Dionex ICS-5000+ ion chromatograph interfaced to an Orbitrap Fusion Tribrid mass spectrometer (Thermo Fisher Scientific, San Jose, CA, USA) operating at a resolution setting of 500,000 (FWHM at m/z 200) on MS1 acquisition to capture all ^13^C isotopologues. The chromatograph was outfitted with a Dionex IonPac AG11-HC-4 µm RFIC&HPIC (2 x 50 mm) guard column upstream of a Dionex IonPac AS11-HC-4 µm RFIC&HPIC (2 x 250 mm) column. Chromatography and mass spectrometric settings were the same as described previously (*55*) with an m/z range of 80 to 700. Metabolites and their isotopologues were identified and their peak areas were integrated and exported to Excel via the TraceFinder 3.3 (Thermo) software package. Peak areas were corrected for natural abundance as previously described(*56*), after which fractional enrichment and µmoles metabolites/g protein were calculated to quantify ^13^C incorporation into various metabolites.

### Online survival analysis

An online survival analysis tool Kaplan Meier plotter(*57*) (http://kmplot.com/analysis/), which includes both clinical data and gene expression data of lung, breast, gastric and ovarian cancers, was used to evaluate the prognostic significance of different genes in NSCLC. According to the median expression value of a gene, the patient samples were divided into high and low expression groups. In this study, the analysis was carried out under the default parameters for histology, stage, grade, gender and smoking history. AJCC stage T=2, N=0 and M=0 was applied. ‘univariate’ for Cox regression and ‘exclude biased arrays’ were utilized for array quality control and in each survival plot, the log rank P-value and hazard ratio (HR, 95% confidence intervals) were calculated and displayed on the main plot.

### Statistical Analysis

Cell viability assay, wound healing assay, cell invasion assay, colony formation, flow cytometry, IF staining, immunohistochemistry (IHC) staining, mitochondrial stress test, qRT-PCR, mitoROS generation, mitochondrial membrane potential, mitophagy assay and tumor mass data were analyzed by unpaired Student t-test. Correlations of PLK1 and PDHA1 were analyzed by linear regression test, and tumor growth for different groups was analyzed by Two-way ANOVA test. All data are presented as the mean±SEM. Data were analyzed using the GraphPad Prism 8 software package. *P<0.05; **P<0.01; ***P<0.001; ****P<0.0001; n.s., not significant.

## Acknowledgements

The research was generously supported by NIH R01 CA157429 (X. Liu), R01 CA196634 (X. Liu), R01 CA264652 (X. Liu), R01 CA256893 (X. Liu). This research was also supported by the Biospecimen Procurement & Translational Pathology, Biostatistics and Bioinformatics, Redox Metabolism, and Flow Cytometry and Immune Monitoring Shared Resources of the University of Kentucky Markey Cancer Center (P30CA177558). Research reported here was supported by an Institutional Development Award (IDeA) from the National Institute of General Medical Sciences of the National Institutes of Health under grant number P30 GM127211. We thank Dr. Richard M. Higashi, Dr. Whei-Mei T. Fan and Dr. Andrew N. Lane for technical assistance and discussion with MS, NMR and related data analysis. We sincerely appreciate the critical reading of the manuscript by Eleanor Erikson and Andrew Lane.

## Competing Interests

The authors declare that they have no competing interest.

---

**Supplementary Figure 1.**
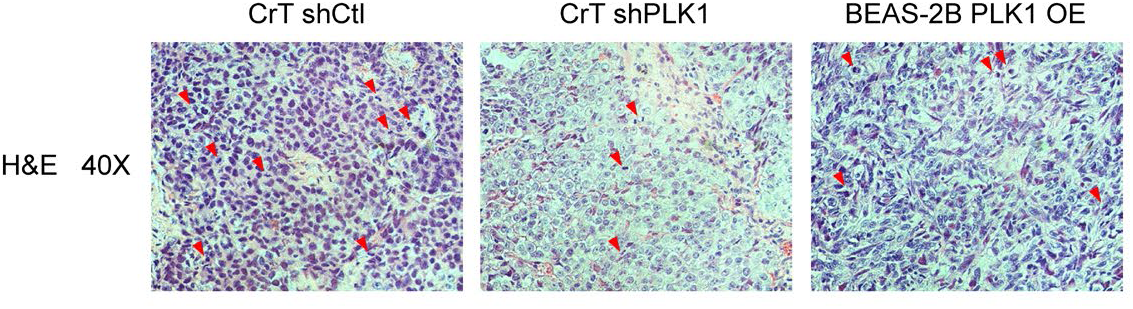
(A) Parental BEAS-2B cells with empty vector and BEAS-2B cells stably overexpressing PLK1 were inoculated into flanks of nude mice (n=15). Image of tumors weight. (B) Representative images of hematoxylin and eosin staining of xenograft tumors derived from CrT shCtl, CrT shPLK1 and BEAS-2B PLK1 OE cells. Red arrows indicate mitotic cells.

**Supplementary Figure 2.**
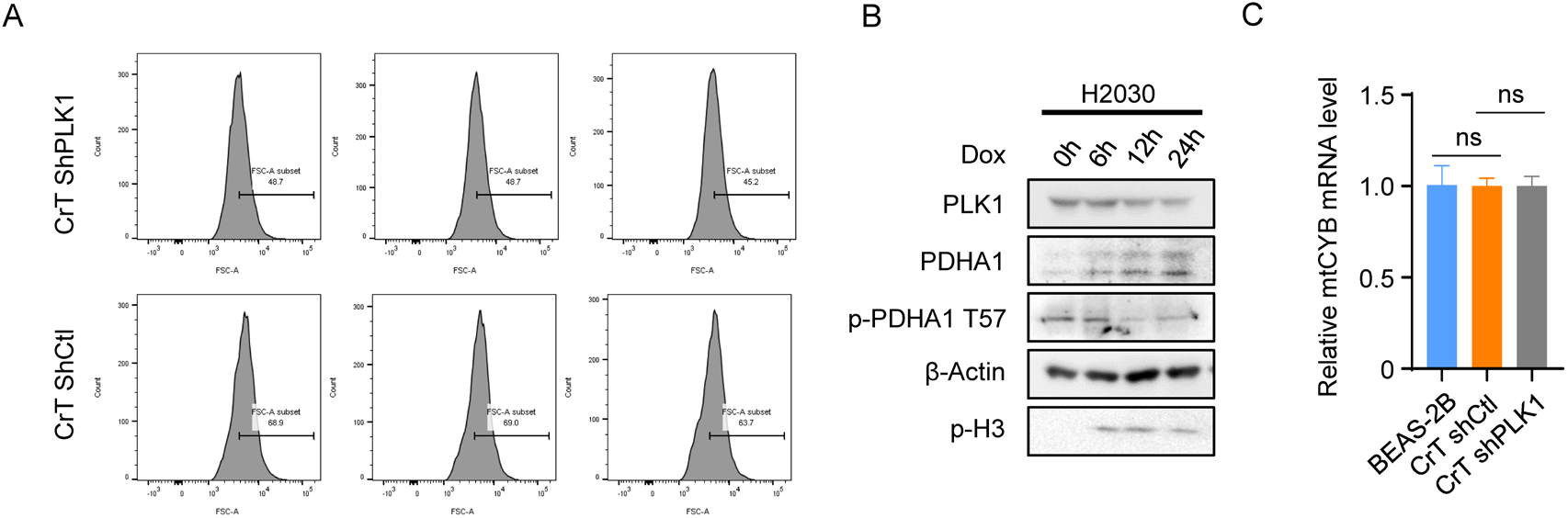
PLK1-associated OXPHOS reprogramming. (A) Flow cytometric analysis of rhodamine 123 in CrT shCtl and CrT shPLK1 cells. (B) IB analysis of protein levels of PLK1 and PDHA1 in Tet-inducible PLK1 knockdown H2030 cells treated with doxycycline for the time indicated. (C) qRT-PCR analysis of the transcriptional level of mtCYB in BEAS-2B, CrT shCtl and CrT shPLK1 cells, followed by quantification, ns P>0.05.

**Supplementary Figure 3.**
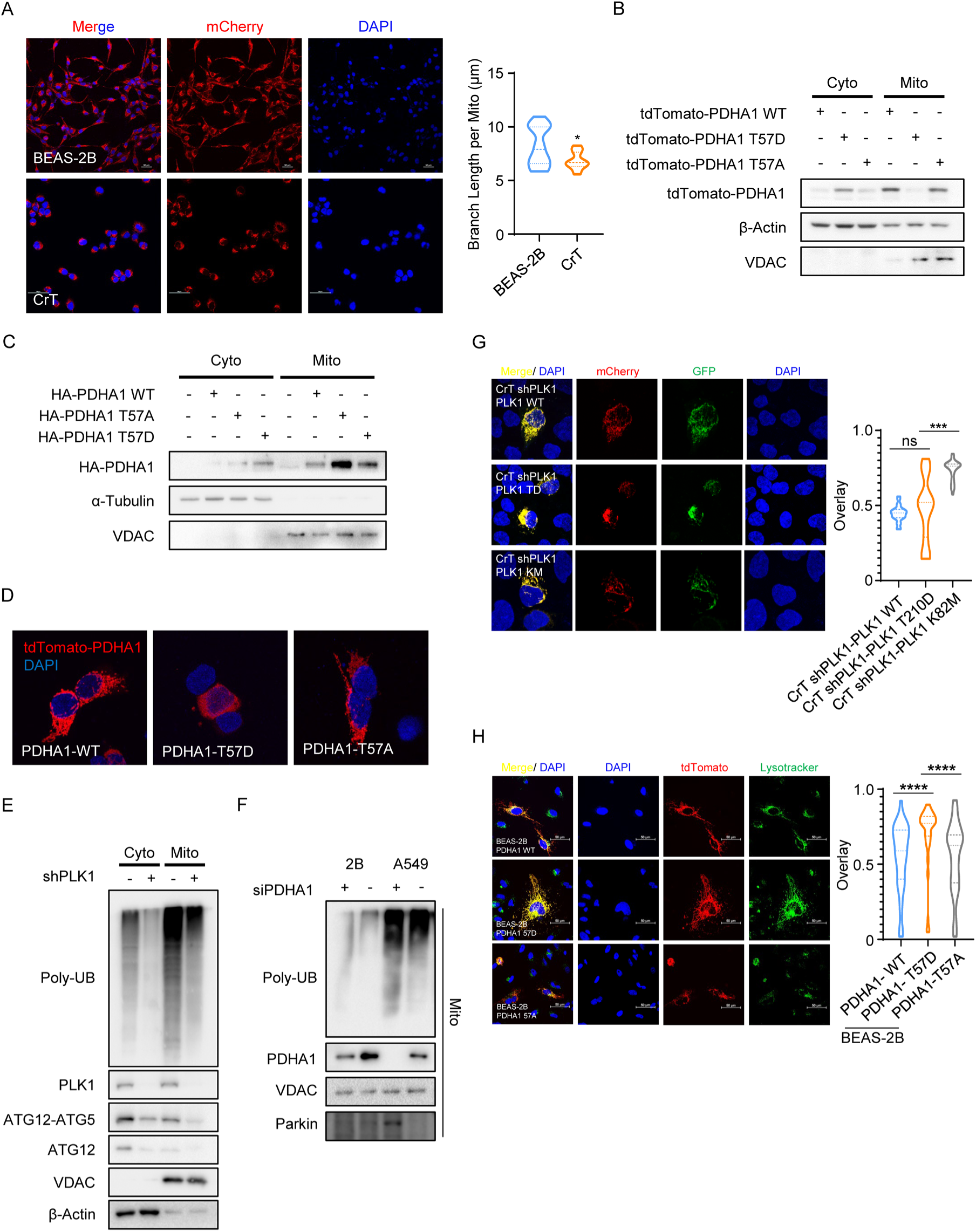
PLK1-associated PDHA1 phosphorylation triggers mitophagy activation and promotes PDHA1 protein degradation. (A) BEAS-2B and CrT cells were incubated with MitoTracker (red), followed by confocal microscopy analysis. After single cells were analyzed using ImageJ plug Marcros, quantitative analysis and comparison of mitochondrial mean branch length between BEAS-2B and CrT cells are presented. *P<0.05. (B) HEK293TA cells were transfected with different forms of PDHA1 constructs (WT, T57D and T57A), and harvested for subcellular fractionation, followed by IB to compare the levels of PDHA1 in cytosol (Cyto) and mitochondria (Mito). (C) IB analysis of protein levels of PDHA1 in cytosol and mitochondria of CrT cells stably expressing different forms of PDHA1 (WT, T57D and T57A). (D) BEAS-2B cells were transfected with different forms of tdTomato-PDHA1 constructs (WT, T57D and T57A) for 2 days, and analyzed by confocal microscopy. Representative images indicate the subcellular localization of PDHA1 variants. (E) CrT shCtl and CrT shPLK1 cells were harvested for subcellular fractionation, followed by IB to address the level of poly-ubiquitination (Poly-UB), PLK1, ATG12 and ATG12-ATG5 conjugation. (F) PDHA1 knockdown by siRNA in BEAS-2B and A549 cells, followed by mitochondrial isolation and IB. (G) CrT shPLK1 cells were transfected with different forms of PLK1 constructs (WT, T210D and K82M), re-transfected with the plasmid expressing the mitophagy indicator for 2 days, and analyzed by the confocal microscopy, followed by quantification of mitophagy activation. ***P<0.001; ns P>0.05. (I) BEAS-2B cells were transfected with different forms of tdTomato-PDHA1 constructs (WT, T57D and T57A) for 2 days, incubated with lysotracker, and analyzed by confocal microscopy, followed by quantification of PDHA1 and lysosome co-localization. ****P<0.0001.

**Supplementary Figure 4.**
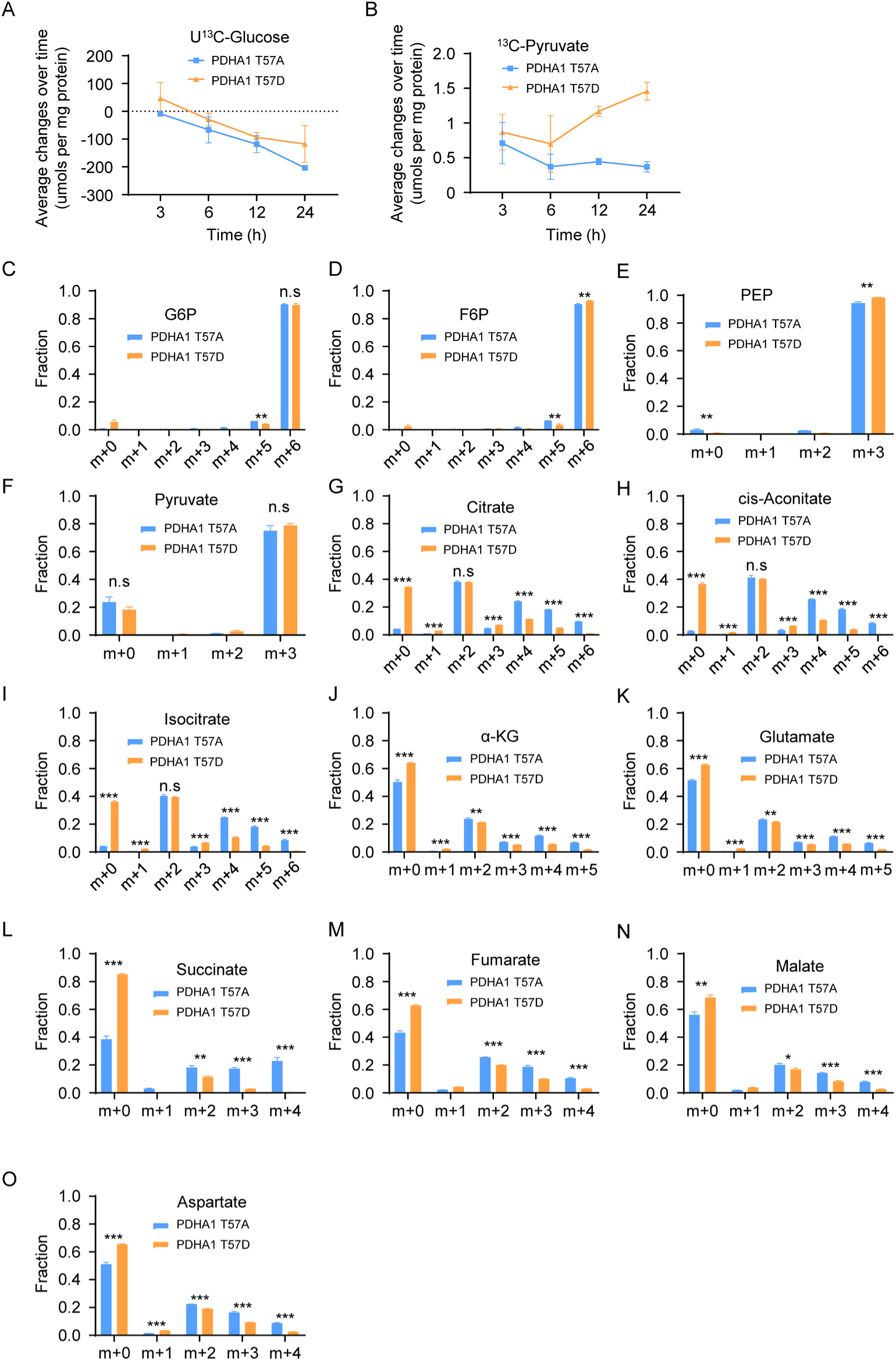
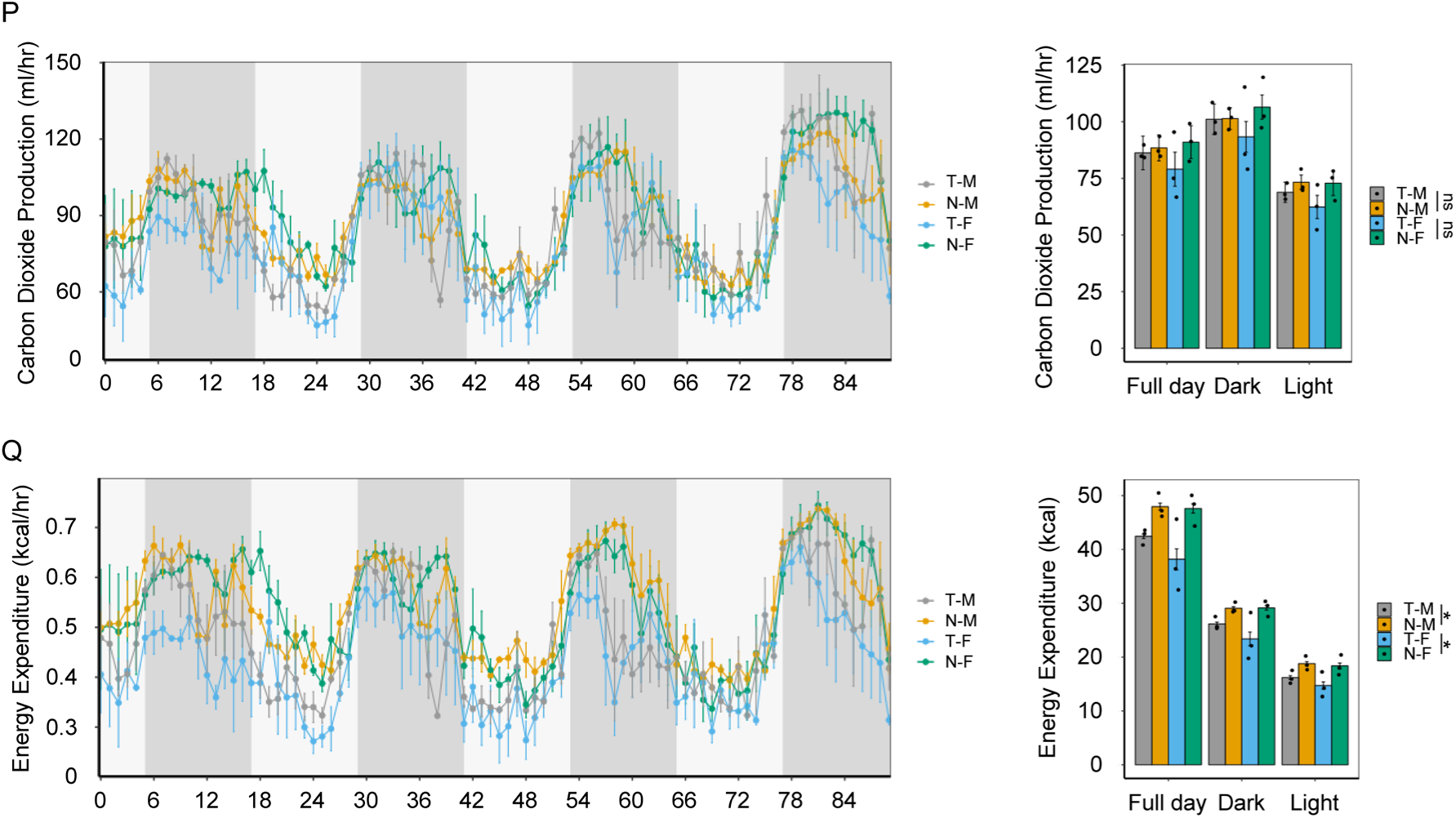
PLK1-mediated PDHA1-T57 phosphorylation affects glucose metabolism and TCA cycle. (A) Time courses of ^13^C enrichment of glucose in media. Representative NMR analysis of ^13^C-enriched glucose in media culturing CrT cells expressing PDHA1 T57A or T57D variant. Media was collected from the plates in 3h, 6h, 12h and 24h and processed to NMR analysis. (B) Representative NMR analysis of ^13^C-enriched pyruvate in media. (C-O) Representative IC-MS analysis of fractional enrichment of ^13^C-labeled central metabolites (glycolysis and TCA cycle) in CrT cells expressing PDHA1 T57A or T57D variant. The x-axis denotes the number of ^13^C atoms present in each compound. The fractional enrichment of isotopologues are presented as means ± standard deviations (n=3), *P<0.05; **P<0.01; ***P<0.001; ****P<0.0001. Carbon dioxide production (P) and energy expenditure (Q) for four groups of mice (T-M, N-M, T-F and N-F) monitored for 84 hours at room temperature. *P<0.05

**Supplementary Figure 5.**
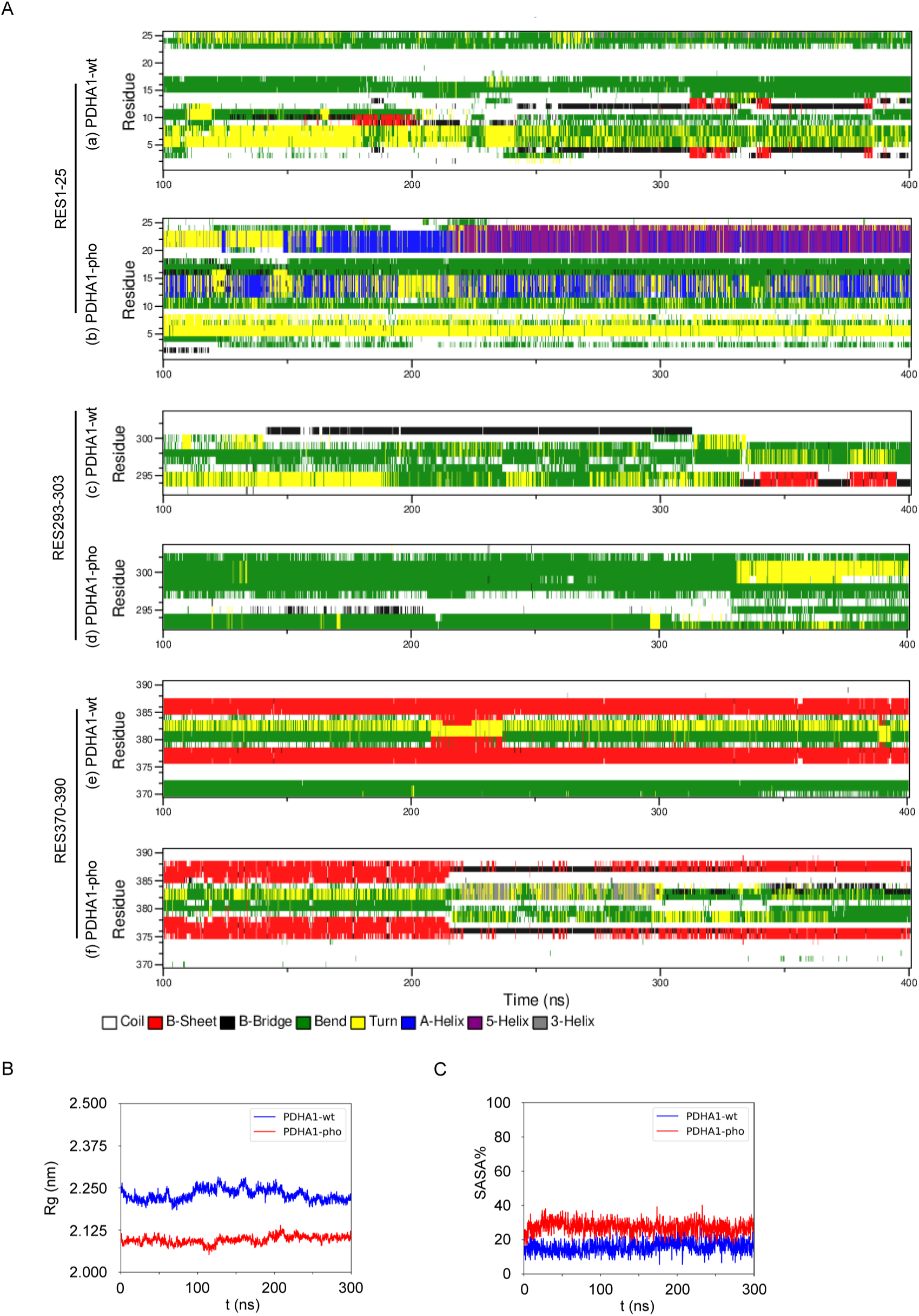
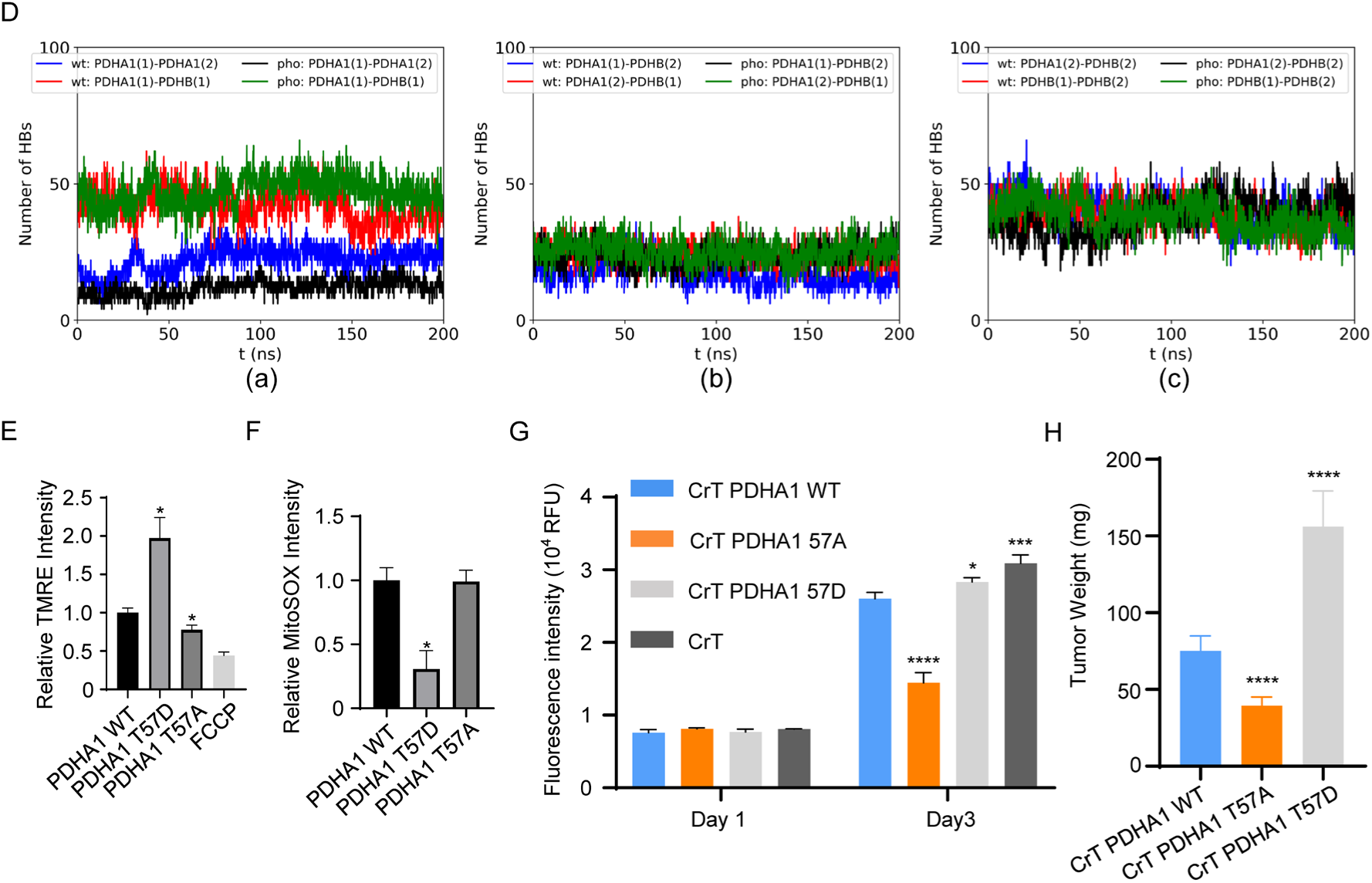
PLK1-mediated PDHA1-T57 phosphorylation downregulates PDHC activity and mitochondrial function to promote tumor growth. (A) Secondary structure of individual residues vs. time for the PDHA1-WT and PDHA1-T57pho in RES1-25 (a and b), RES293-303 (c and d) and RES370-390 (e and f). (B) The radius of gyration of PDHA1-WT and PDHA1-T57pho protein vs. time. (C) The SASA% of PDHA1-WT and PDHA1-T57pho protein vs. time. (D) The average HBs vs. time in PDHA1-wt-PDHB and PDHA1-pho-PDHB heterotetramers. (E) Quantification of mitochondrial ROS level in HEK293TA cells transiently expressing PDHA1 variants (WT, T57D and T57A) is presented as means ± standard deviations (n=8). *P<0.05. (F) Quantification of mitochondrial membrane potential level in HEK293TA cells transiently expressing PDHA1 variants (WT, T57D and T57A) is presented as means ± standard deviations (n=12).*P<0.05. (G) MTT analyses of CrT cells stably expressing wild-type PDHA1 and PDHA1 variants (T57D and T57A) are presented as means ± standard deviations (n=6). *P<0.05; ***P<0.001; ****P<0.0001. (H) Tumor growth of CrT cells stably expressing wild-type PDHA1 and PDHA1 variants (T57D and T57A) in nude mice (n=6). tumor weight is presented. ****P<0.0001.

**Supplementary Figure 6.**
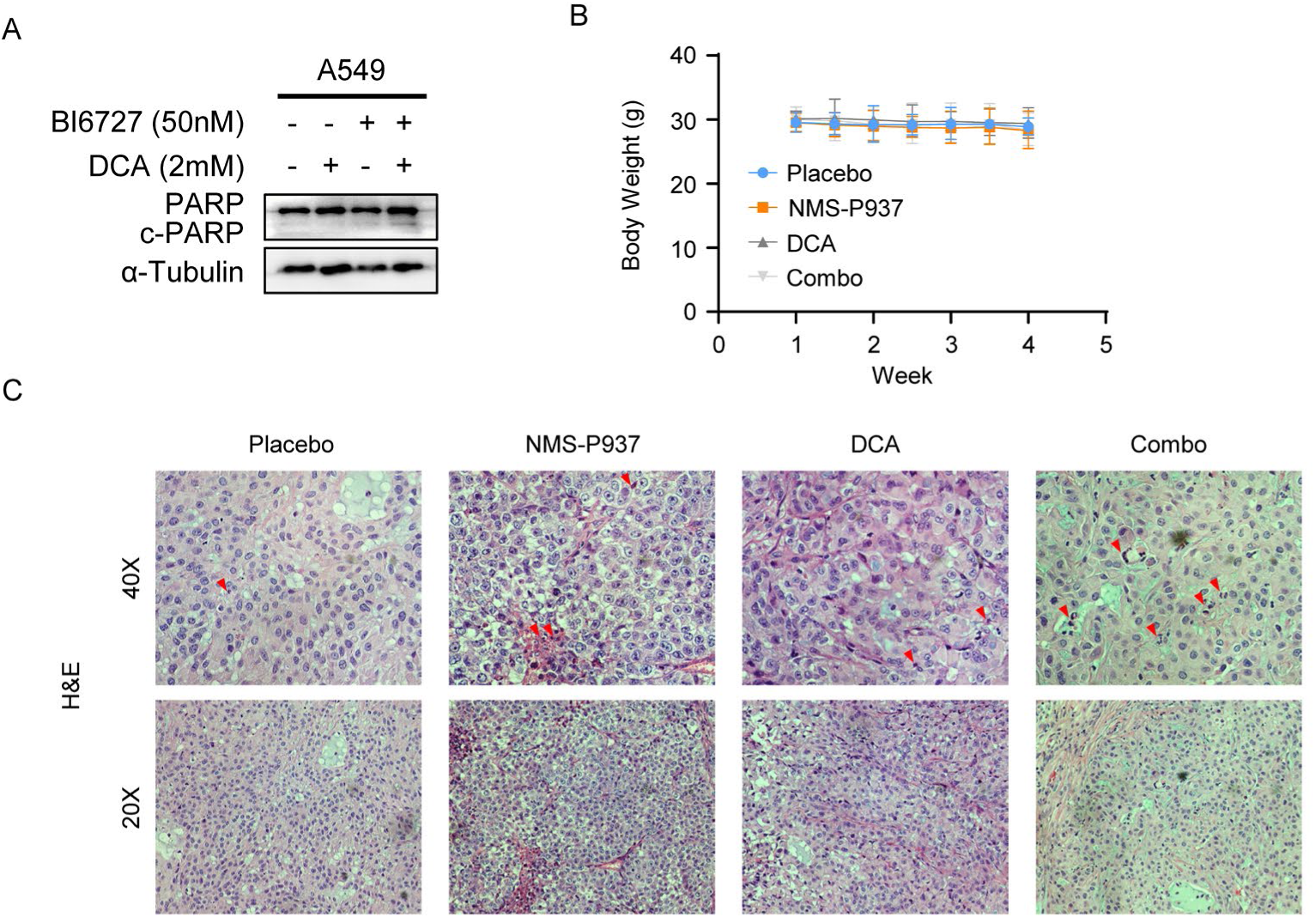
PLK1 inhibition enhances chemo-sensitivity to DCA in vitro and in vivo. (A) A549 cells were treated with 50nM BI6727 and 2mM DCA as indicated, followed by IB analysis of c-PARP protein level. (B) Body weights of mice during the treatment period. (C) Effects of DCA and NMS-P937 on tumor growth. Representative images of hematoxylin and eosin staining of xenograft tumors derived from CrT cells. Red arrows indicate apoptotic cells.

